# Recognition of discrete export signals in early flagellar subunits during bacterial Type III secretion

**DOI:** 10.1101/2020.12.09.414946

**Authors:** Owain J. Bryant, Paraminder Dhillon, Colin Hughes, Gillian M. Fraser

## Abstract

Type III Secretion Systems (T3SS) deliver subunits from the bacterial cytosol to nascent cell surface flagella. Early flagellar subunits that form the rod and hook substructures are unchaperoned and contain their own export signals. A gate recognition motif (GRM) docks them at the FlhBc component of the FlhAB-FliPQR export gate, but the gate must then be opened and subunits must be unfolded to pass through the flagellar channel. This induced us to seek further signals on the subunits. Here, we identify a second signal at the extreme N-terminus of flagellar rod and hook subunits and determine that key to the signal is its hydrophobicity. We show that the two export signal elements are recognised separately and sequentially, as the N-terminal signal is recognised by the flagellar export machinery only after subunits have docked at FlhB_C_ *via* the GRM. The position of the N-terminal hydrophobic signal in the subunit sequence relative to the GRM appeared to be important, as a FlgD deletion variant (FlgD_short_), in which the distance between the N-terminal signal and the GRM was shortened, ‘stalled’ at the export machinery and was not exported. The attenuation of motility caused by FlgD_short_ was suppressed by mutations that destabilised the closed conformation of the FlhAB-FliPQR export gate, suggesting that the hydrophobic N-terminal signal might trigger opening of the flagellar export gate.

## Introduction

Type III Secretion Systems (T3SS) are multi-component molecular machines that deliver protein cargo from the bacterial cytosol either to their site of assembly in cell surface flagella or virulence factor injectisomes, or directly to their site of action in eukaryotic target cells or the extracellular environment [1–5]. The flagellar T3SS (fT3SS) directs the export of thousands of structural subunits required for the assembly and operation of flagella, rotary nanomotors for cell motility that extend from the bacterial cell surface [1, 6]. Newly synthesised subunits of the flagellar rod, hook and filament are targeted to the fT3SS, where they are unfolded and translocated across the cell membrane, powered by the proton motive force and ATP hydrolysis, into an external export channel that spans the length of the nascent flagellum [7], [8]. During flagellum biogenesis, when the rod/hook structure reaches its mature length, the fT3SS switches export specificity from recognition of ‘early’ rod/hook subunits to ‘late’ subunits for filament assembly [9, 10]. This means that early and late flagellar subunits must be differentiated by the fT3SS machinery to ensure that they are exported at the correct stage of flagellum biogenesis. This is achieved, in part, by targeting subunits to the export machinery at the right time using a combination of export signals in the subunit mRNA and/or polypeptide. T3SS substrates contain N-terminal signals for targeting to the export machinery, however they do not share a common peptide sequence [11–15]. In addition, some substrates are piloted to the T3SS machinery by specific chaperones [16–20].

The core export components of the fT3SS are evolutionarily related to those of the virulence injectisome, with which they share considerable structural and amino acid sequence similarity [22–25]. The flagellar export machinery comprises an ATPase complex (FliHIJ) located in the cytoplasm, peripheral to the membrane. Immediately above the ATPase is a nonameric ring formed by the cytoplasmic domain of FlhA (FlhA_C_), which functions as a subunit docking platform [20, 21, 26, 27]. A recent cryo-ET map indicates that the FlhA family have a sea-horse-like structure, in which FlhAc forms the ‘body’ and the FlhA N-terminal region (FlhA_N_) forms the ‘head’, which is fixed in the plane of the membrane [28]. FlhA_N_ wraps around the base of a complex formed by FliPQR and the N-terminal sub-domain of FlhB (FlhB_N_), and together these form the FlhAB-FliPQR export gate that connects the cytoplasm to the central channel in the nascent flagellum, which is contiguous with the extracellular environment [22–29]. FlhB_N_ is connected *via* a linker (FlhB_CN_) to the cytoplasmic domain of FlhB (FlhB_C_), which is thought to sit between the FlhA_N_ and FlhA_C_ rings, where it functions as a docking site for early flagellar subunits [14, 22, 28].

The ‘early’ flagellar subunits that assemble to form the rod and hook substructures are not chaperoned: instead, the signals for targeting and export are found within the early subunits themselves. We have shown that of one of these signals is a small hydrophobic sequence termed the gate recognition motif (GRM), which is essential for early subunit export [14]. This motif binds a surface exposed hydrophobic pocket on FlhBc [14]. Once subunits reach the export machinery, they must be unfolded before they can pass through the narrow channel formed by FliPQR-FlhB_N_ into the central channel of the nascent flagellum, through which the subunits transit until they reach the tip and fold into the structure [6, 22]. Structural studies suggest that FliPQR-FlhB_N_ adopts an energetically favourable closed conformation, possibly to maintain the membrane permeability barrier [22, 25, 30, 31]. This suggests that there must be a mechanism to trigger opening of the export gate when subunits dock at cytosolic face of the flagellar export machinery.

Here we sought to identify new export signals within flagellar rod/hook subunits, using the hook-cap subunit FlgD as a model export substrate. We show that the extreme N-terminus of rod/hook subunits contains a hydrophobic export signal and investigate its functional relationship to the subunit gate recognition motif (GRM).

## Results

### Identification of a hydrophobic export signal at the N-terminus of FlgD

The N-terminal region of flagellar rod and hook subunits is required for their export [12, 14]. Using the flagellar hook-cap protein FlgD as a model rod/hook subunit, we sought to identify specific export signals within the N-terminus. A screen of ten FlgD variants containing internal five-residue scanning deletions in the first 50 residues (though FlgDΔ2-5 is a four-residue deletion, retaining the initial methionine) identified just two variants defective for export into culture supernatant (Fig. 1A). Loss of residues 2-5 caused a significant reduction in export, as did deletion of residues 36-40, though to a lesser extent (Fig. 1A; [14]). We have shown that FlgD residues 36-40 are the gate recognition motif (GRM) required for transient subunit docking at the FlhB_C_ component of the export gate [14]. The results suggest that the extreme N-terminus might also be important for interaction with the export machinery.

**Figure 1.**
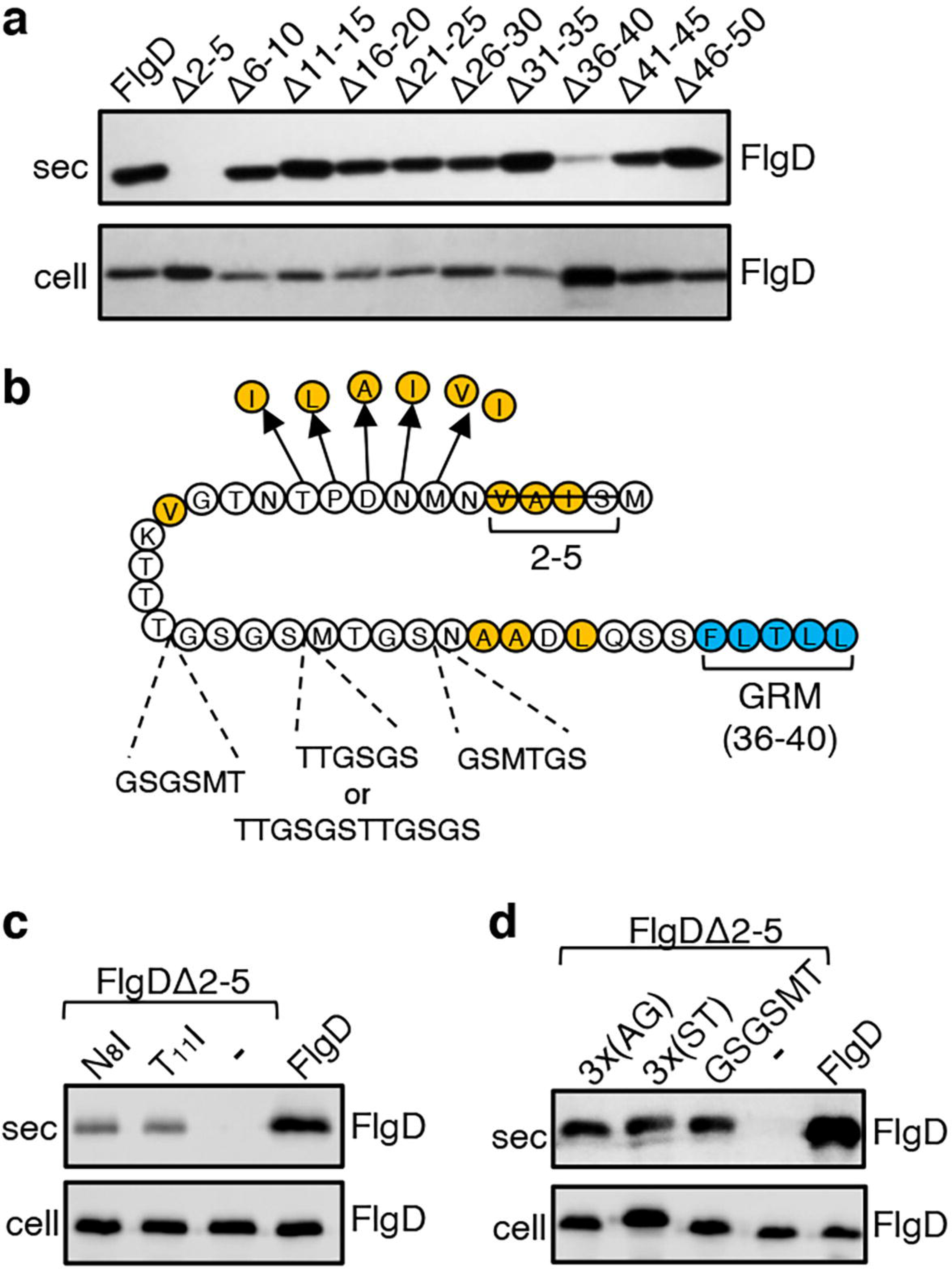
Screening for export-defective FlgD variants. **a.** Whole cell (cell) and supernatant (sec) proteins from late exponential phase cultures of a *Salmonella flgD* null strain expressing plasmid-encoded wild type FlgD (FlgD) or its variants (Δ2-5, Δ6-10, Δ11-15, Δ16-20, Δ21-25, Δ26-30, Δ31-35, Δ36-40, Δ41-45 or Δ46-50) were separated by SDS (15%)-PAGE and analysed by immunoblotting with anti-FlgD polyclonal antisera. **b.** A schematic displaying all intragenic suppressor mutations within amino acids 1-40 of FlgD isolated from the FlgDΔ2-5 variant. Small non-polar residues are highlighted in orange. All suppressor mutations were located between the gate-recognition motif (GRM, blue) and the extreme N-terminus, and can be separated into two classes: insertions or duplications that introduced additional sequence between valine-15 and the gate-recognition motif, or missense mutations that re-introduce small non-polar residues at the N-terminus. All intragenic suppressors isolated from FlgDΔ2-5 are displayed in this figure. **c.** Whole cell (cell) and supernatant (sec) proteins from late exponential-phase cultures of *Salmonella flgD* null strains expressing plasmid-encoded: suppressor mutants isolated from the FlgDΔ2-5 variant (FlgDΔ2-5-N_8_I or FlgDΔ2-5-T_11_I), FlgDΔ2-5 variant (-) or wild type FlgD (FlgD) were separated by SDS (15%)-PAGE and analysed by immunoblotting with anti-FlgD polyclonal antisera. **d.** Whole cell (cell) and supernatant (sec) proteins from late exponential phase cultures of *Salmonella flgD* null strains expressing plasmid-encoded wild type FlgD (labelled FlgD), FlgDΔ2-5 (labelled as DΔ2-5) or variants of FlgDΔ2-5 containing between residues 19 and 20 a six-residue insertion of either small non-polar (AGAGAG) residues (labelled as 3x(AG)), polar (STSTST) residues (labelled as 3x(ST)), or the sequence from an isolated insertion suppressor mutant (GSGSMT) (labelled as GSGSMT), were separated by SDS (15%)-PAGE and analysed by immunoblotting with anti-FlgD polyclonal antisera.

To gain insight into the putative new signal, we screened for intragenic suppressor mutations that could restore export of the FlgDΔ2-5 variant. A *Salmonella flgD* null strain expressing *flgD*Δ2-5 *in trans* was inoculated into soft-tryptone agar and incubated until ‘spurs’ of motile cell populations appeared. Sequencing of *flgD*Δ2-5 alleles from these motile populations identified ten different intragenic gain-of-function mutations. These could be separated into two classes (Fig. 1B**, Fig. S1**).

The first class of motile revertants carried *flgD*Δ2-5 alleles with missense mutations that introduced small non-polar residues at the extreme N-terminus of FlgDΔ2-5 (Fig. 1B). Deletion of residues 2-5 (^2^SIAV^5^) had removed all small non-polar amino acids from the first ten residue region of FlgD, effectively creating a new N-terminus containing a combination of polar, charged or large non-polar residues (**Fig. S2**). Analysis of other flagellar rod and hook subunit primary sequences revealed that in every case their native N-terminal regions contain small non-polar residues positioned upstream of the gate recognition motif (GRM residues 36-40; **Fig. S2**), indicating that hydrophobicity may be key to the function of the N-terminal export signal. Removal of non-polar residues from the extreme N-terminus of the secreted hook-length control protein, FliK, attenuated its export, indicating that the hydrophobic N-terminal signal is required for export of early subunits (**Fig. S3**).

Export assays performed with two representative motile revertant strains carrying *flgDΔ2-5* variants with gain-of-function point mutations, those encoding FlgDΔ2-5-N_8_I and FlgDΔ2-5-T_11_I, revealed that export of these subunits had recovered to ∼50% of the level observed for wild type FlgD (Fig. 1C**, Fig. S4-S5**).

The second class of motile revertants carried *flgD*Δ2-5 alleles that had acquired duplications or insertions introducing at least six additional residues between the FlgDΔ2-5 N-terminus and the gate recognition motif (GRM; Fig. 1B). It seemed possible that these insertions/duplications might have restored subunit export either by insertion of amino acids that could function as a ‘new’ hydrophobic export signal, or by restoring the position of an existing small hydrophobic residue or sequence relative to the GRM.

To assess these possibilities, we tested whether export of FlgDΔ2-5 could be recovered by inserting either polar (^19^STSTST^20^) or small non-polar (^19^AGAGAG^20^) residues in the FlgDΔ2-5 N-terminal region at a position equivalent to one of the suppressing duplications (^19^GSGSMT^20^; Fig. 1B **and** D**, Fig. S4-S5**). We reasoned that if suppression by the additional sequence had been caused by repositioning an existing small hydrophobic amino acid relative to the GRM, then any insertional sequence (polar or non-polar) would restore export, while if suppression had resulted from insertion of a ‘new’ export signal, then either the polar STSTST or non-polar AGAGAG, but not both, could be expected to restore export.

We found that both the engineered FlgD variants (FlgDΔ2-5-^19^AGAGAG^20^ and FlgDΔ2-5-^19^STSTST^20^) were exported from a *Salmonella flgD* null strain as effectively as the gain-of-function mutant FlgDΔ2-5-^19^GSGSMT^20^ isolated from the suppressor screen (Fig. 1D). This suggests that the insertions had repositioned a sequence in the FlgDΔ2-5 N-terminus relative to the GRM to overcome the loss of small hydrophobic residues.

### The position of the hydrophobic export signal relative to the gate recognition motif is critical for rod and hook subunit export

The intragenic suppressor experiments indicated that FlgD export requires a hydrophobic export signal towards the N-terminus and that the position of this hydrophobic signal relative to the previously described GRM is important. Sequence analysis of the gain-of-function FlgDΔ2-5 insertion variants revealed that the insertions were all located between the GRM and valine_15_ (V_15_; Fig. 1B). We reasoned that they repositioned valine_15_ relative to the GRM, such that it could perform the function of the N-terminal hydrophobic signal lost in FlgDΔ2-5. To test this view, we replaced V_15_ by alanine in the gain-of-function variant FlgDΔ2-5-^19^(GSGSMT)^20^ and assayed its export in the *Salmonella flgD* null (Fig. 2A**, Fig. S6**). Unlike the *flgD* null strain producing either the parental FlgDΔ2-5-^19^(GSGSMT)^20^ or wild type FlgD, the *flgD* null carrying variant FlgDΔ2-5-^19^(GSGSMT)^20^-V_15_A was non-motile, reflecting the variant’s failure to export (Fig. 2A**, Fig. S6**). This suggests that the V_15_ residue had indeed compensated for the missing N-terminal hydrophobic signal.

**Figure 2.**
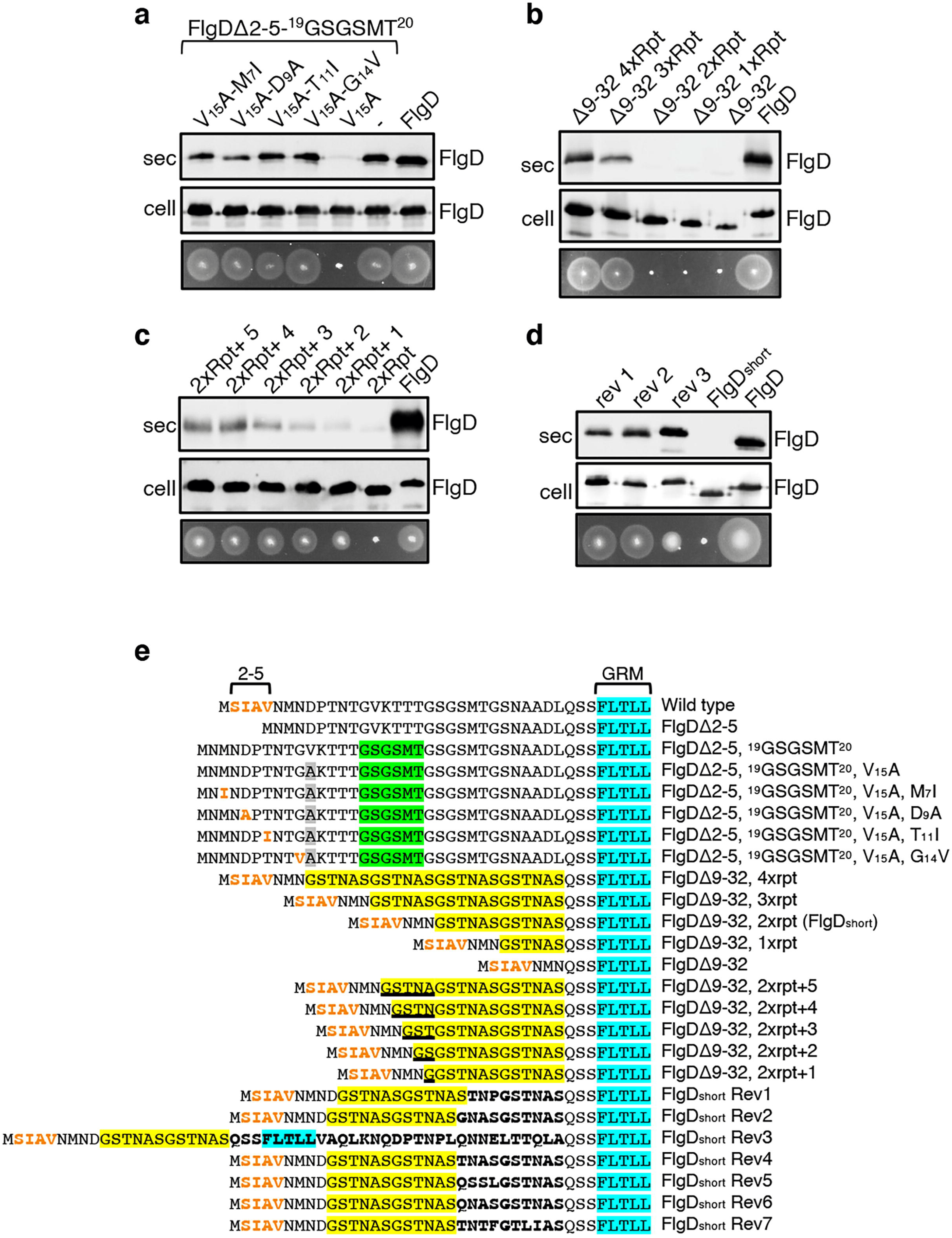
Export of FlgD variants in which the position of the hydrophobic export signal is varied relative to the gate recognition motif (GRM). Whole cell (cell) and supernatant (sec) proteins from late exponential-phase cultures of a *Salmonella flgD* null strain expressing plasmid encoded suppressor mutants isolated from the FlgDΔ2-5-^19^(GSGSMT)^20^-V_15_A variant (V_15_A-M_7_I, V_15_A-D_9_A, V_15_A-T_11_I, V_15_A-G_14_V), their parent FlgD variant FlgDΔ2-5-^19^(GSGSMT)^20^-V_15_A (labelled as V_15_A), FlgDΔ2-5-^19^(GSGSMT)^20^ (labelled as -) or wild type FlgD (FlgD) were separated by SDS (15%)-PAGE and analysed by immunoblotting with anti-FlgD polyclonal antisera. Swimming motility (bottom panel; 0.25% soft tryptone agar) of the same strains were carried out at 37°C for 4-6 hours. **b.** Whole cell (cell) and supernatant (sec) proteins from late exponential-phase cultures of a *Salmonella flgD* null strain expressing plasmid-encoded wild type FlgD (labelled as FlgD), FlgDΔ9-32 or its variants in which residues 9-32 were replaced by between one and four six-residue repeats of Gly-Ser-Thr-Asn-Ala-Ser (GSTNAS): (Δ9-32 4xRpt, Δ9-32 3xRpt, Δ9-32 2xRpt or Δ9-32 1xRpt) were separated by SDS (15%)-PAGE and analysed by immunoblotting with anti-FlgD polyclonal antisera. Swimming motility (bottom panel; 0.25% soft tryptone agar) of the same strains were carried out at 37°C for 4-6 hours. **c.** Whole cell (cell) and supernatant (sec) proteins from late exponential-phase cultures of a *Salmonella flgD* null strain expressing plasmid-encoded wild type FlgD (labelled as FlgD), a FlgD variant in which residues 9-32 were replaced by two repeats of a six-residue sequence Gly-Ser-Thr-Asn-Ala-Ser (labelled as 2xRpt) or its variants containing between one and five additional residues inserted directly after the two repeats (labelled as 2xRpt+1, 2xRpt+2, 2xRpt+3, 2xRpt+4 or 2xRpt+5) were separated by SDS (15%)-PAGE and analysed by immunoblotting with anti-FlgD polyclonal antisera. Swimming motility (bottom panel; 0.25% soft tryptone agar) of the same strains were carried out at 37°C for 4-6 hours. **d.** Whole cell (cell) and supernatant (sec) proteins from late exponential-phase cultures of a *Salmonella flgD* null strain expressing plasmid-encoded wild type FlgD (labelled as FlgD), a FlgD variant in which residues 9-32 were replaced by two repeats of a six-residue sequence Gly-Ser-Thr-Asn-Ala-Ser (labelled as FlgD_short_) or suppressor mutants isolated from this strain (labelled as rev1, rev2 or rev3) were separated by SDS (15%)-PAGE and analysed by immunoblotting with anti-FlgD polyclonal antisera. Swimming motility (bottom panel; 0.25% soft tryptone agar) of the same strains were carried out at 37°C for 4-6 hours. **e.** N-terminal sequences of wild type FlgD and its variants aligned to their gate-recognition motif (GRM; blue). The following sequence features or residues are displayed: The N-terminal hydrophobic signal (residue 2-5; orange), the Gly-Ser-Gly-Ser-Met-Thr (GSGSMT) insertion (green) isolated from the FlgDΔ2-5 suppressor screen, the valine-15 to alanine mutation (grey), small non-polar mutations (M7I, D9A, T11I, G14V; orange) isolated from the FlgDΔ2-5, ^19^GSGSMT^20^ suppressor screen, FlgDΔ9-32 and its variants in which residues 9-32 are replaced with one, two, three or four repeats of a six-residue sequence Gly-Ser-Thr-Asn-Ala-Ser (GSTNAS; yellow), FlgDΔ9-32, 2xrpt (hereafter termed FlgD_short_) containing five, four, three, two, or one additional residues (underlined) inserted between the GRM and N-terminal hydrophobic signal, and suppressor mutants (Rev 1-7) isolated from FlgD_short_ that introduced additional residues (bold) between the N-terminal hydrophobic signal (orange) and the gate-recognition motif (blue).

By screening for intragenic suppressors of the motility defect associated with FlgDΔ2-5-^19^(GSGSMT)^20^-V_15_A, four gain-of-function missense mutations were identified, M_7_I, D_9_A, T_11_I and G_14_V. All of these had introduced small hydrophobic residues, all positioned at least 27 residues upstream of the GRM. These FlgDΔ2-5-^19^(GSGSMT)^20^-V_15_A gain-of-function variants restored motility to the *Salmonella flgD* null strain and were exported at levels similar to wildtype FlgD and FlgDΔ2-5-^19^(GSGSMT)^20^ (Fig. 2A**, Fig. S6**). These data confirm the importance of small non-polar residues positioned upstream of the GRM.

Our results so far had indicated that the position of the FlgD N-terminal hydrophobic export signal relative to the GRM was critical and suggested that, for export to occur efficiently, at least 26 residues must separate the hydrophobic signal and the GRM (Fig 1C**, Fig. S4**). In the primary sequences of all *Salmonella* flagellar rod/hook subunits the GRM is positioned ≥ 30 amino acids downstream of the subunit N-terminus (**Fig S4**), suggesting that separation of the two signals by a minimum number of residues might be a common feature among early flagellar subunits. To test this, a suite of engineered *flgD* alleles was constructed that encoded FlgD variants in which wildtype residues 9-32 were replaced with between one and four repeats of the six amino acid sequence Gly-Ser-Thr-Asn-Ala-Ser (GSTNAS). Swimming motility and export assays revealed that the minimum number of inserted GSTNAS repeats that could support efficient FlgD export was three, equivalent to separation of the hydrophobic N-terminal signal and the GRM by 24 residues (Fig. 2B). Below this threshold, FlgD export and swimming motility were strongly attenuated (Fig. 2B). A further set of recombinant *flgD* alleles was constructed, which encoded FlgDΔ9-32 variants carrying two GSTNAS repeats (hereafter termed FlgD_short_) directly followed by between one and five additional residues (Fig. 2E). Motility and FlgD export increased incrementally with the addition of each amino acid (Fig. 2C**, Fig. S6**). The data indicate that a low level of FlgD export is supported when the hydrophobic N-terminal signal (_2_SIAV_5_) and the GRM (_36_FLTLL_40_) are separated by 19 residues, with export efficiency and swimming motility increasing as separation of the export signals approaches an optimal 24 residues.

To further establish the requirement for a minimum number of residues between the hydrophobic N-terminal signal and the GRM, we screened for intragenic suppressor mutations that could restore swimming motility in a *flgD* null strain producing FlgD_short_. Sequencing of *flgD_short_* alleles from motile revertant strains identified 7 gain-of-function mutations that introduced additional residues between the hydrophobic N-terminal signal and the GRM (**Fig. S6**). Swimming motility and FlgD export was assessed for three *flgD* null strains expressing representative *flgD_short_* gain-of-function variants and all showed increased FlgD subunit export and swimming motility compared to the *flgD* null expressing *flgD_short_* (Fig. 2D**, Fig. S6**). The data confirm that the position of the hydrophobic N-terminal signal relative to the GRM is critical for efficient FlgD subunit export.

To establish that this is a general requirement for the export of other rod and hook subunits, engineered alleles of *flgE* (hook) and *flgG* (rod) were constructed that encoded variants in which FlgE residues 9-32 or FlgG residues 11-35 were either deleted (FlgE_short_ and FlgG_short_) or replaced with four repeats of the sequence GSTNAS (Fig. 3). As had been observed for FlgD_short_, export of the FlgE_short_ and FlgG_short_ variants was severely attenuated compared to wild type FlgE and FlgG (Fig. 3**, Fig. S7**). Furthermore, insertion of four GSTNAS repeats into FlgE_short_ and FlgG_short_ recovered subunit export to wild type levels, indicating that the minimum separation of the hydrophobic N-terminal signal and the GRM is a feature throughout rod and hook subunits (Fig. 3).

**Figure 3.**
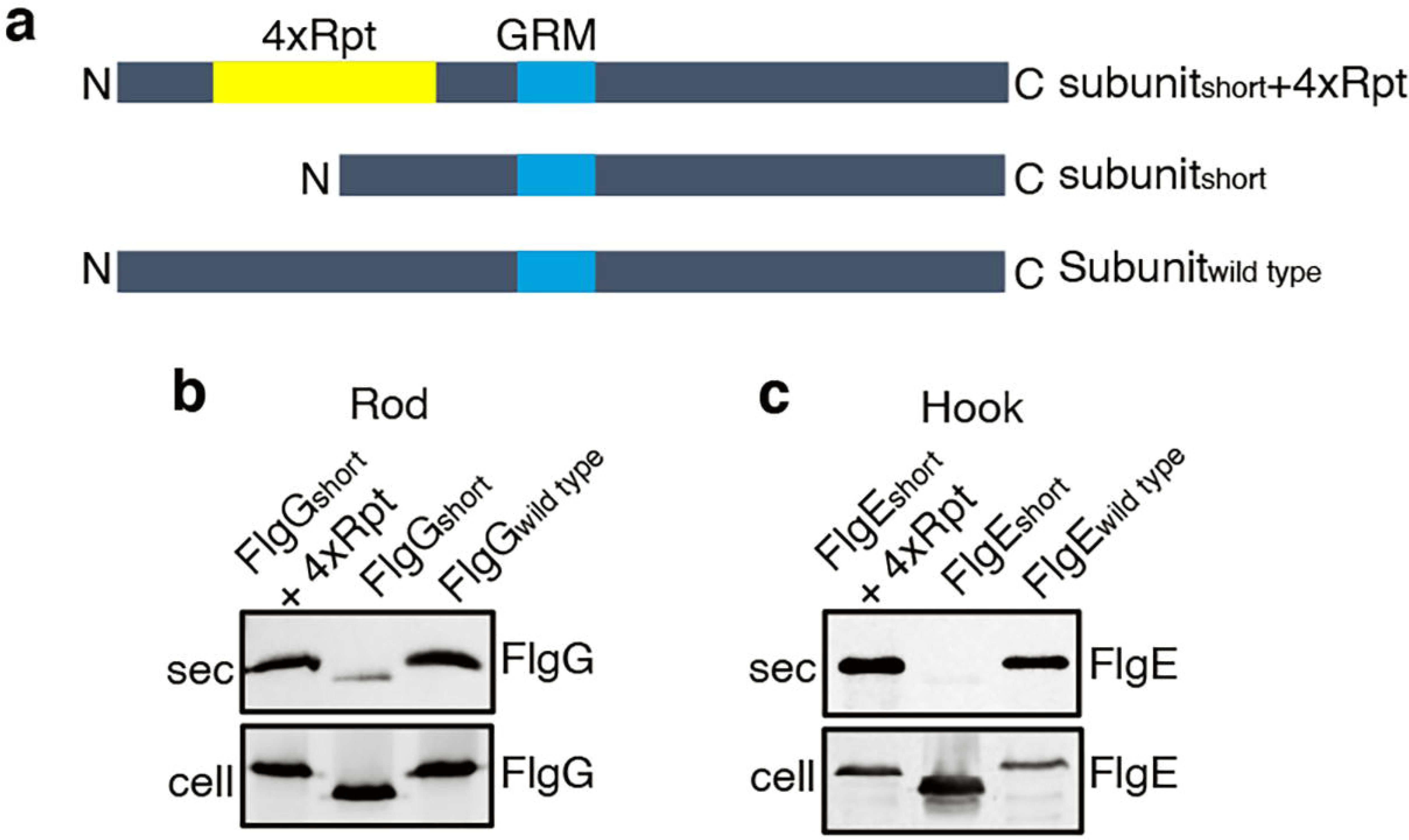
Effect of the relative position of the N-terminus and GRM on the export of other rod and hook subunits. **a.** Schematic representation of a wild type subunit (labelled as subunit_wild type_), a subunit containing a deletion of sequence from between the N-terminus and GRM (labelled as subunit_short_) and a subunit in which the deleted sequence was replaced by four repeats of a six-residue sequence Gly-Ser-Thr-Asn-Ala-Ser (yellow, labelled as subunit_short_+4Rpt). **b.** Whole cell (cell) and supernatant (sec) proteins from late exponential-phase cultures of a *Salmonella flgE* null strain expressing plasmid-encoded wild type FlgG (labelled as FlgG_wild type_), a FlgG variant in which residues 11-35 were deleted (labelled as FlgG_short_) or a FlgG variant in which residues 11-35 were replaced by four repeats of a six-residue sequence Gly-Ser-Thr-Asn-Ala-Ser (labelled as FlgG_short_+4Rpt). All FlgG variants were engineered to contain an internal 3xFLAG tag for immunodetection. Proteins were separated by SDS (15%)-PAGE and analysed by immunoblotting with anti-FLAG monoclonal antisera. **c.** Whole cell (cell) and supernatant (sec) proteins from late exponential-phase cultures of a *Salmonella flgD* null strain expressing plasmid-encoded wild type FlgE (labelled as FlgE_wild type_), a FlgE variant in which residues 9-32 were deleted (labelled as FlgE_short_) or a FlgE variant in which residues 9-32 were replaced by four repeats of a six-residue sequence Gly-Ser-Thr-Asn-Ala-Ser (labelled as FlgE_short_+4Rpt). All FlgE variants were engineered to contain an internal 3xFLAG tag for immunodetection. Proteins were separated by SDS (15%)-PAGE and analysed by immunoblotting with anti-FLAG monoclonal antisera.

### Sequential engagement of the subunit GRM and hydrophobic N-terminal export signal by the flagellar export machinery

Having identified a new hydrophobic N-terminal export signal and established that its position relative to the GRM was critical, we next wanted to determine the order in which the signals were recognised/engaged by the export machinery. The signals might be recognised simultaneously, with both being required for initial entry of rod/hook subunits into the export pathway. Alternatively, they might be recognised sequentially. If this were the case, then a subunit variant that possessed the ‘first’ signal but was deleted for the ‘second’ signal might enter the export pathway but fail to progress, becoming stalled at a specific step to block the pathway and prevent export of wild type subunits. To test if FlgDΔ2-5 or FlgDΔGRM stalled in the export pathway, recombinant expression vectors encoding these variants or wild type FlgD were introduced into a *Salmonella* Δ*recA* strain that is wild type for flagellar export (Fig. 4). We could then assess whether the variant FlgD constructs could interfere *in trans* with the wildtype flagellar export. To do this we assessed the export of an early flagellar substrate, FliK, which controls the length of the flagellar hook and of the late export substrate, FlgK and FlgL, which together form a junction connecting the flagellar filament to the hook. We saw that FlgDΔ2-5 inhibited motility and export of the FliK, FlgK and FlgL flagellar subunits, whereas FlgDΔGRM did not (Fig. 4B **and** C). The data indicate that FlgDΔ2-5 enters the flagellar export pathway and stalls at a critical point, blocking export. In contrast, FlgDΔGRM does not stall or block export.

**Figure 4.**
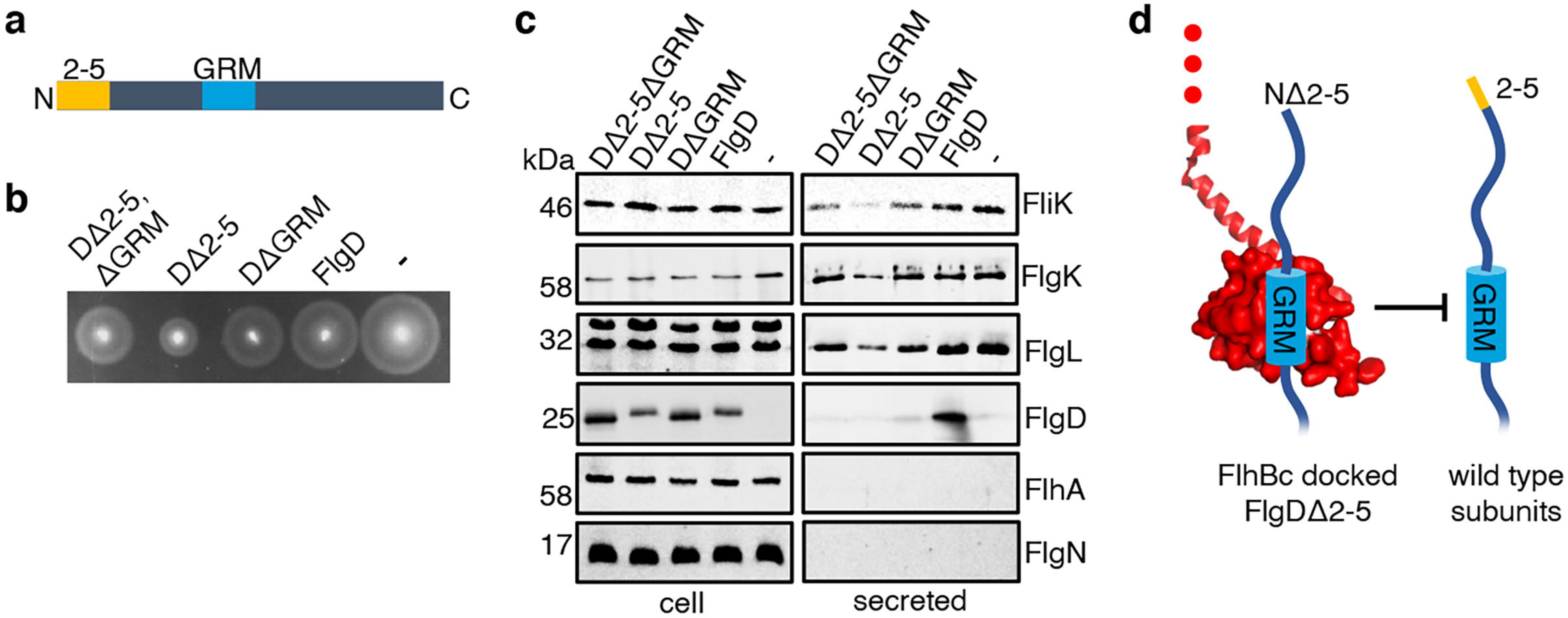
Effect on subunit export of overexpressed FlgDΔ2-5 and variants. **a.** Schematic representation of a FlgD subunit containing a N-terminal hydrophobic signal (orange, labelled as 2-5) and a gate-recognition motif (blue, labelled as GRM). **b.** Swimming motility of a *Salmonella* Δ*recA* strain expressing plasmid-encoded wild type FlgD (FlgD), its variants (DΔ2-5ΔGRM, DΔ2-5 or DΔGRM) or empty pTrc99a vector (-). Motility was assessed in 0.25% soft-tryptone agar containing 100 μg/ml ampicillin and 100μM IPTG and incubated for 4-6 hours at 37°C. **c.** Whole cell (cell) and secreted proteins (secreted) from late-exponential-phase cultures were separated by SDS (15%)-PAGE and analysed by immunoblotting with anti-FliK (hook ruler subunit), anti-FlgK and anti-FlgL (hook-filament junction subunits), anti-FlgD (hook cap subunit, anti-FlhA (component of the export machinery) and anti-FlgN (export chaperone for FlgK and FlgL) polyclonal antisera. Apparent molecular weights are in kilodaltons (kDa). **d.** A model depicting a FlgDΔ2-5 subunit (left) docked *via* its gate-recognition motif (GRM, blue) at the subunit binding pocket on FlhB_C_ (PDB: 3B0Z[31], red), preventing wild type subunits (right) from docking at FlhB_C_.

To determine whether FlgDΔ2-5 stalls at a point before or after subunit docking at the FlhB_C_ component of the flagellar export gate *via* the GRM, a recombinant vector encoding a FlgD variant in which both export signals were deleted (FlgDΔ2-5ΔGRM) was constructed. If loss of the hydrophobic N-terminal signal had caused subunits to stall after docking at FlhB_C_, then additional deletion of the subunit GRM would relieve this block. Motility and subunit export assays revealed that the *Salmonella* Δ*recA* strain producing FlgDΔ2-5ΔGRM displayed swimming motility and levels of FliK and FlgK subunit export similar to cells producing FlgDΔGRM (Fig. 4B **and** C). The data suggest that FlgDΔ2-5 stalls after docking at the FlhB_C_ export gate, preventing docking of other early subunits.

It seemed possible that subunit docking *via* the GRM to the FlhB_C_ export gate might position the hydrophobic N-terminal signal in close proximity to its recognition site on the export machinery. If this were the case, ‘short’ subunit variants containing deletions that decreased the number of residues between the hydrophobic N-terminal signal and the GRM might also stall at FlhB_C_, and this stalling might be relieved by additional deletion of the GRM. To test this, recombinant expression vectors encoding ‘short’ subunit variants (FlgE_short_ or FlgD_short_), ‘short’ subunit variants additionally deleted for the GRM (FlgE_short_ΔGRM or FlgD_short_ΔGRM) or wild type FlgE or FlgD were introduced into a *Salmonella* Δ*recA* strain (Fig 5). Compared to the wild type subunits expressed *in trans*, the ‘short’ subunits inhibited swimming motility and the export of other flagellar subunits (FliD, FliK, FlgK), whereas FlgE_short_ΔGRM and FlgD_short_ΔGRM did not (Fig 5). Taken together with the data presented in Figure 4, the results indicate that the subunit GRM and the hydrophobic N-terminal signal are recognised sequentially, with subunits first docking at FlhB_C_ *via* the GRM, which positions the hydrophobic N-terminal signal for subsequent interactions with the export machinery (Fig 4D).

**Figure 5.**
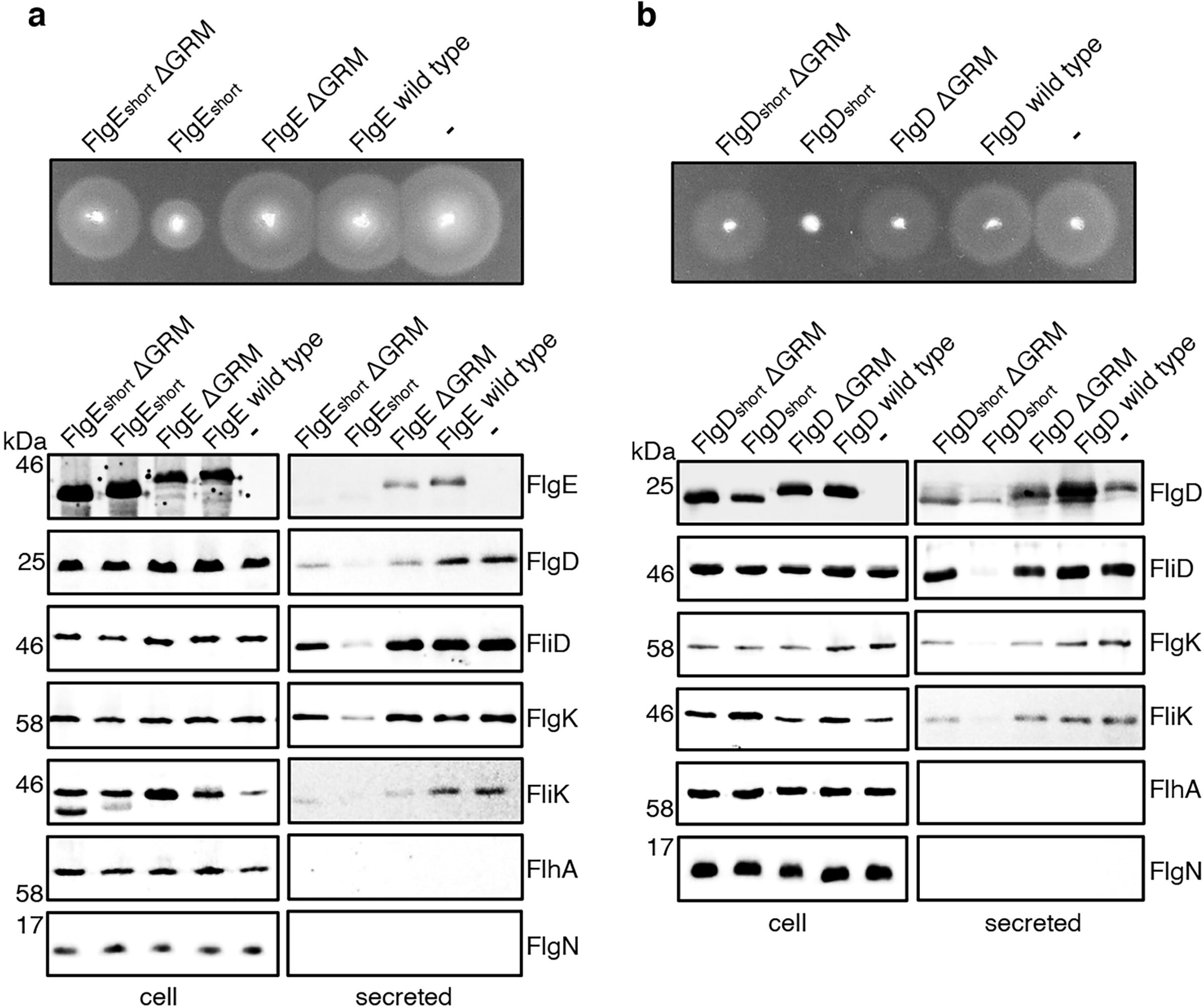
Effect on subunit export of overexpressed FlgE_short,_ FlgD_short_ and variants. **a.** Swimming motility of a *Salmonella* Δ*recA* strain expressing plasmid-encoded wild type FlgE (labelled as FlgE wild type), a FlgE variant in which residues 9-32 were deleted (labelled as FlgE_short_), a FlgE variant in which residues 9-32 and residues 39-43 (corresponding to the gate-recognition motif) were deleted (labelled as FlgE_short_ΔGRM), a FlgE variant in which residues 39-43 were deleted (labelled as FlgEΔGRM) or empty pTrc99a vector (labelled as -). All FlgE variants were engineered to contain an internal 3xFLAG tag for immunodetection. Motility was assessed in 0.25% soft-tryptone agar containing 100 μg/ml ampicillin and 100 μM IPTG and incubated for 4-6 hours at 37°C (top panel). Whole cell (cell) and secreted proteins (secreted) from late-exponential-phase cultures were separated by SDS (15%)-PAGE and analysed by immunoblotting with anti-FLAG monoclonal antisera (to detect the flag tagged hook subunit FlgE) or anti-FlgD (hook cap subunit), anti-FliD (filament cap subunit), anti-FlgK (hook-filament junction subunit), anti-FliK (hook-ruler subunit), anti-FlhA (component of the export machinery) and anti-FlgN (chaperone for FlgK and FlgL) polyclonal antisera (bottom). Apparent molecular weights are in kilodaltons (kDa). **b.** Swimming motility of a *Salmonella* Δ*recA* strain expressing plasmid-encoded wild type FlgD (labelled as FlgD wild type), a FlgD variant in which residues 9-32 were replaced with two repeats of the six amino acid sequence Gly-Ser-Thr-Asn-Ala-Ser (labelled as FlgD_short_), a FlgD variant in which residues 9-32 were replaced with two repeats of the six amino acid sequence Gly-Ser-Thr-Asn-Ala-Ser and residues 36-40 were deleted (labelled FlgD_short_ΔGRM), a FlgD variant in which residues 36-40 were deleted (labelled as FlgDΔGRM) or empty pTrc99a vector (labelled as -). Motility was assessed in 0.25% soft-tryptone agar containing 100 μg/ml ampicillin and 100 μM IPTG and incubated for 4-6 hours at 37°C (top panel). Whole cell (cell) and secreted proteins (secreted) from late-exponential-phase cultures were separated by SDS (15%)-PAGE and analysed by immunoblotting with anti-FlgD (hook cap subunit), anti-FliD (filament cap subunit), anti-FlgK (hook-filament junction subunit), anti-FliK (hook ruler subunit), anti-FlhA (component of export machinery) and anti-FlgN (chaperone for FlgK and FlgL) polyclonal antisera (bottom). Apparent molecular weights are in kilodaltons (kDa).

### Mutations that promote opening of the export gate partially compensate for incorrect positioning of the N-terminal export signal

The accruing data indicated that subunit docking at FlhB_C_ might correctly position the hydrophobic N-terminal signal for recognition by the export machinery. To model the position of FlhB_C_ relative to other components of the export machinery, we docked the structures of FlhB_C_ and the FliPQR-FlhB_N_ export gate into the tomographic reconstruction of the *Salmonella* SPI-1 type III secretion system (Fig 6B) [32]. The model indicated that FliPQR-FlhB_N_ and the subunit docking site on FlhB_C_ are separated by a minimum distance of ∼78 Å, and that FlhB_C_ is positioned no more than ∼22-45 Å from FlhA (Fig 6A; [33]). Taking FlgD as a model early flagellar subunit, the distance between the FlgD N-terminal hydrophobic signal and the GRM was found to be in the range of ∼45 Å (*α*-helix) to ∼105 Å (unfolded contour length), depending on the predicted structure adopted by the subunit N-terminus (Fig 6A). Based on these estimates, it seemed feasible that the hydrophobic N-terminal signal of a subunit docked at FlhB_C_ could contact either FlhA or the FliPQR-FlhB_N_ complex, and that this interaction might trigger opening of the export gate [22, 32, 34, 35]. If this were true, mutations that promote the open conformation of the export gate might compensate for the incorrect positioning of the hydrophobic N-terminal export signal in ‘short’ rod/hook subunits. One such export gate mutation, FliP-M_210_A, has been shown to increase ion conductance across the bacterial inner membrane, indicating that this gate variant fails to close efficiently [31].

**Figure 6.**
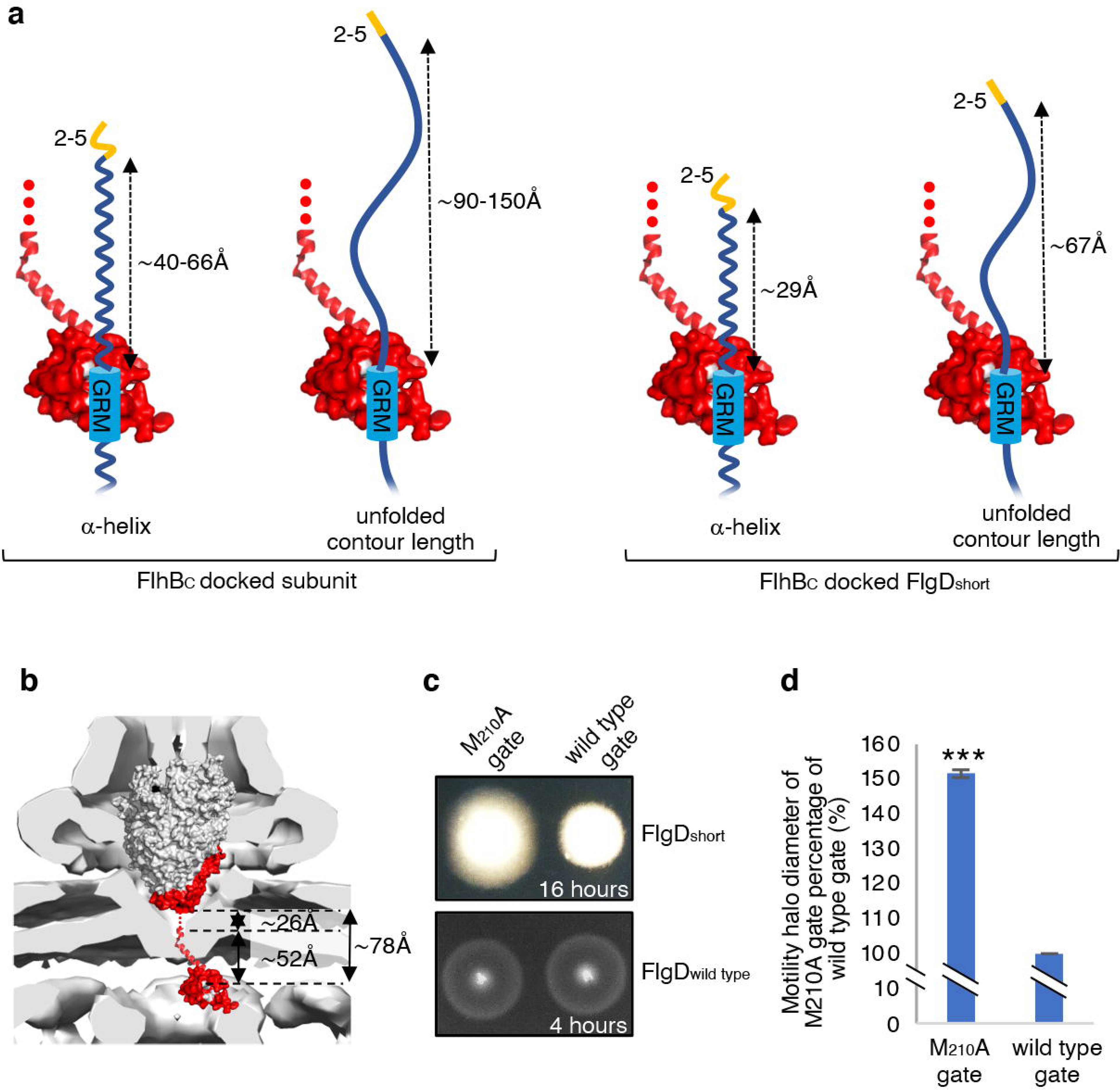
Suppression of the FlgD_short_ motility defect by mutations in FliP. **a.** A model depicting subunits docked *via* their gate-recognition motif (GRM, blue) at the subunit binding pocket on FlhB_C_ (PDB: 3B0Z [32], red) with N-termini of early flagellar subunits adopting either an *α*-helical conformation separating the N-terminal hydrophobic signal (2-5, orange) and gate-recognition motif (GRM, blue) by ∼40-60 ångstrom (where each amino acid is on average separated by ∼1.5Å, left) or an unfolded conformation where the unfolded contour length separating the N-terminal hydrophobic signal (2-5) and gate-recognition motif (GRM) is ∼90-150 ångstrom (where each amino acid is on average separated by ∼3.5Å, middle left). Values corresponding to the distance separating the N-terminal hydrophobic signal (2-5, orange) and gate-recognition motif (GRM, blue) of a FlgD subunit variant in which residues 9-32 are replaced with two repeats of the six amino acid sequence Gly-Ser-Thr-Asn-Ala-Ser (FlgD_short_) indicate that the N-terminal hydrophobic signal (2-5, orange) and gate-recognition motif (GRM, blue) are separated by ∼29 ångstrom (*α*-helical conformation, middle right) or ∼67 ångstrom (unfolded contour length, right). **b.** Placement of the crystal structure of FlhBc (PDB:3BOZ [31]; red) and the cryo-EM structure of FliPQR-FlhB (PDB:6S3L[30]) in a tomographic reconstruction of the *Salmonella* SPI-1 injectisome (EMD-8544 [60]; grey). The minimum distance between the subunit gate-recognition motif binding site on FlhB_C_ (grey) to FlhB_N_ (defined as *Salmonella* FlhB residue 211 [61]; ∼78Å) was estimated by combining: the value corresponding to the distance between the subunit binding pocket on FlhB_C_ [14] (grey) and the N-terminal visible residue (D_229_) in the FlhB_C_ structure (PDB:3BOZ [31]; ∼52Å) with the value corresponding to the minimum distance between FlhB residues 211 and 228 (based on a linear *α*-helical conformation; ∼26Å). **c.** Swimming motility of recombinant *Salmonella flgD* null strains producing a chromosomally-encoded FliP-M_210_A variant (M_210_A gate, left) or wild type FliP (wild type gate, right). Wild type FliP and FliP-M_210_A were engineered to contain an internal HA tag positioned between residue 21 and 22 to allow immunodetection of FliP. Both strains produced either a pTrc99a plasmid-encoded FlgD subunit variant in which residues 9-32 were replaced with two repeats of the six amino acid sequence Gly-Ser-Thr-Asn-Ala-Ser (FlgD_short_; top panel) or a pTrc99a plasmid-encoded wild type FlgD subunit (FlgD_wild type_; bottom panel). Motility was assessed in 0.25% soft-tryptone agar containing 100 μg/ml ampicillin and 50 μM IPTG and incubated for 16 hours (top panel) or 4-6 hours at 37°C (bottom panel). **d.** The mean motility halo diameter of recombinant *Salmonella flgD* null strains producing a chromosomally-encoded FliP-M_210_A variant (M_210_A gate, left) or wild type FliP (wild type gate, right). Wild type FliP and FliP-M_210_A were engineered to contain an internal HA tag positioned between residue 21 and 22 to allow immunodetection of FliP. Both strains produced a pTrc99a plasmid-encoded FlgD subunit variant in which residues 9-32 were replaced with two repeats of the six amino acid sequence Gly-Ser-Thr-Asn-Ala-Ser (FlgD_short_). Error bars represent the standard error of the mean calculated from at least three biological replicates. *** indicates a p-value <0.001.

To test whether the FliP-M_210_A variant gate could promote export of ‘short’ subunits, in which the distance between the hydrophobic N-terminal signal and the GRM was reduced, a recombinant expression vector encoding FlgD_short_ was introduced into *Salmonella flgD* null strains in which the *fliP* gene had been replaced with recombinant genes encoding either a functional FliP variant with an internal HA-tag (designated wild type gate) or the equivalent HA-tagged FliP-M_210_A variant (designated M_210_A gate; Fig. 6C**, Fig. S8**). The swimming motility of these strains was found to be consistently stronger in the strain producing the M_210_A gate compared to the strain with the wild type gate, with the motility halo of the *fliP*-M210A-Δ*flgD* strain expressing FlgD_short_ having a 50% greater diameter than that of the wild type *fliP*-Δ*flgD* strain expressing FlgD_short_ (Fig. 6D**, Fig. S8**). This increase in motility indicated that the defect caused by incorrect positioning of the hydrophobic N-terminal signal relative to the GRM in FlgD_short_ could indeed be partially compensated by promoting the gate open conformation.

## Discussion

T3SS substrates contain N-terminal export signals, but these have not been fully defined and how they promote subunit export remains unclear. Here, we characterised a new hydrophobic N-terminal export signal in early flagellar rod/hook subunits and showed that the position of this signal relative to the known subunit gate recognition motif (GRM) is key to subunit export.

Loss of the hydrophobic N-terminal signal in the hook cap subunit FlgD had a stronger negative effect on subunit export than deletion of the GRM that enables subunit docking at FlhB_C_, suggesting that the hydrophobic N-terminal signal may be required to trigger an essential export step. A suppressor screen showed that the export defect caused by deleting the hydrophobic N-terminal signal could be overcome by mutations that either reintroduced small non-polar amino acids positioned 3-7 residues from the subunit N-terminus (*e.g.* M_7_I), or introduced additional residues between V_15_ and the GRM. In such ‘gain of function’ strains containing insertions, changing V_15_ to alanine abolished subunit export, which was rescued by re-introduction of small non-polar residues close to the N-terminus. These data point to an essential export function for small non-polar residues close to the N-terminus of rod/hook subunits.

It was fortuitous that we chose FlgD as the model for early flagellar subunit. All early subunits contain small hydrophobic residues close to the N-terminus, but FlgD is unique in that only four (I_3_, A_4_, V_5_ and V_15_) of its first 25 residues are small and non-polar (**Fig. S1**). Indeed, there are only three other small hydrophobic residues between the FlgD N-terminus and the GRM (Fig. 1**, S1, S9**). While deletion of residues 2-5 in FlgD removes the critical hydrophobic N-terminal signal, similar deletions in the N-terminal regions of other rod/hook subunits reposition existing small non-polar residues close to the N-terminus (**Fig. S1, S9**). This is perhaps why previous deletion studies in early flagellar subunits have failed to identify the hydrophobic N-terminal signal [14, 36, 37].

The finding that subunits lacking the hydrophobic N-terminal export signal, but not the GRM, stalled during export suggested that these two signals were recognised by the flagellar export machinery in a specific order. Mutant variants of other flagellar export substrates or export components have been observed to block the export pathway. For example, a FlgN chaperone variant lacking the C-terminal 20 residues stalls at the FliI ATPase [18], while a GST-tagged FliJ binds FlhA but is unable to associate correctly with FliI so blocking wild type FliJ interaction with FlhA [38]. These attenuations can be reversed by further mutations that disrupt the stalling interactions. This was also observed for FlgDΔ2-5. Loss of the hydrophobic N-terminal signal resulted in a dominant-negative effect on motility and flagellar export, but this was abolished by subsequent deletion of the GRM. This indicates that FlgDΔ2-5 stalls in the export pathway at FlhB_C_, blocking the binding site for early flagellar subunits. These data are consistent with sequential recognition of the two export signals: the GRM first docking subunits at FlhB_C_, and positioning the hydrophobic N-terminal signal to trigger the next export step.

The position of the subunit hydrophobic N-terminal export signal relative to the GRM appears critical for export. Engineering of *flgD* to encode variants in which the region between the N-terminus and the GRM was replaced with polypeptide sequences of varying lengths showed that these signals must be separated by a minimum of 19 residues for detectable export, with substantial export requiring separation by 30 residues (Fig. 3). When subunits dock at FlhB_C,_ which is likely situated within or just below the plane of the inner membrane, the hydrophobic N-terminal signal is positioned close to the FlhAB-FliPQR export gate (Fig. 6B). Subunits in which the GRM and N-terminal signal are brought closer together stall at FlhB_C_, suggesting that the hydrophobic N-terminal signal is unable to contact its recognition site on the export machinery (Figs. 5, 6A **and** 6B). In all flagellar rod/hook subunits, the GRM is positioned at least 30 residues from the N-terminus (**Fig. S1, S9**). The physical distance between the two signals will depend on the structure adopted by the subunit N-terminus (Fig. 6A). The N-terminal region of flagellar subunits is often unstructured in solution [11–15, 35], and such disorder may be an intrinsic feature of flagellar export signals [11-15, 39, 40], as they are typical in other bacterial export N-terminal substrate signals such as those of the Sec and Tat systems [41–42]. Unstructured signals may facilitate multiple interactions with different binding partners during export, and in the case of export systems that transport unfolded proteins they may aid initial entry of substrates into narrow export channels [41].

As yet, nothing is known about the structure of the subunit N-terminal domain upon interaction with the flagellar export machinery. Signal peptides in TAT pathway substrates switch between disordered and α-helical conformations depending on the hydrophobicity of the environment [43]. It therefore seems likely that local environments along the flagellar export pathway will influence the conformation of subunit export signals [22, 44]. Interestingly, FlgD variants that contain additional sequence that position the N-terminal export signal and GRM further apart far above the required threshold did not impede export, which argues that the sequence between both export signals is unfolded (Fig. 2).

If the region between the hydrophobic N-terminal signal and the GRM is unstructured and extended, this would correspond to a polypeptide contour length of approximately 72-105 Å (where the length of one amino acid is ∼3.6 Å). If the same region were to fold as an α-helix, its length would be approximately 30-36 Å (where one amino acid rises every ∼1.5 Å). Placement of the AlphaFold predicted structure of full length FlhB into a tomogram of the T3SS suggests that FlhB_C_ is positioned below the plane of the inner membrane but above the nonameric ring of FlhA_C_ (**Fig. S10**). Without further structural information on subunit interactions with the flagellar export machinery and the precise position of FlhB_C_ within the machinery, it is difficult to determine precisely where the hydrophobic N-terminal signal contacts the machinery.

We speculate that one function of the subunit hydrophobic N-terminal signal might be to trigger opening of the FlhAB-FliPQR export gate, which rests in an energetically favourable closed conformation to maintain the permeability barrier across the bacterial inner membrane [22, 28, 30, 31, 34]. The atomic resolution structure of FliPQR showed that it contains three gating regions [22]. FliR provides a loop (the R-plug) that sits within the core of the structure. Below this, five copies of FliP each provides three methionine residues that together form a methionine-rich ring (M-gasket) under which ionic interactions between adjacent FliQ subunits hold the base of the structure shut (Q-latch). Mutational and evolutionary analyses have shown that the R-plug, M-gasket and Q-latch stabilize the closed export gate conformation to maintain the membrane barrier, preventing the leakage of small molecules whilst allowing the passage of substrates into the export channel [34, 45].

We have also shown that the FliPQR export gate opens and closes in response to export substrate availability, indicating that the export gate reverts to a closed conformation in the absence of export substrates, thereby maintaining the integrity of the cell membrane [34]. These data indicate that the export gate must be opened in response to substrate docking at the export machinery [34, 45].

If the function of the subunit hydrophobic N-terminal signal was to trigger opening of this gate, we hypothesised that mutations which destabilised the gate’s closed conformation would suppress the motility defect associated with FlgD_short_, in which the distance between the N-terminal signal and the GRM is reduced. Introduction of the FliP-M_210_A mutation, which partially destabilises the gate’s closed state, did indeed partly suppress the FlgD_short_ motility defect. We did not find export gate variants that completely destabilised gate closure, but it may be that such mutations disrupt the membrane permeability barrier [30]. This could also explain why in screens for suppressors of FlgDΔ2-5 or FlgD_short_ we did not isolate mutations in genes encoding export gate components (data not shown).

The surface-exposed hydrophobic GRM-binding pocket on FlhB_C_ is well conserved across the T3SS SctU family, to which FlhB belongs [14, 46, 47]. Furthermore, the GRM is conserved in all four injectisome early subunits (SctI, SctF, SctP, OrgC) and is located at least 30 residues away from small hydrophobic residues near the subunit N-terminus (**Fig. S11**). It therefore seems plausible that the ‘dual signal’ mechanism we propose for early flagellar export operates in all T3SS pathways.

In many other pathways, the presence of a substrate triggers opening or assembly of the export channel. The outer membrane chitin transporter in *Vibrio* adopts a closed conformation in which the N-terminus of a neighbouring subunit acts as a pore plug [48]. Chitin binding to the transporter ejects the plug, opening the transport channel and allowing chitin transport [48]. In the Sec pathway, interactions of SecA, ribosomes or pre-proteins with SecYEG can induce conformational changes that promote channel opening [41, 49–51]. In the TAT system, which transports folded substrates across the cytoplasmic membrane, substrate binding to the TatBC complex triggers association with, and subsequent polymerisation of, TatA, which is required for substrate translocation [42, 52]. All of these mechanisms serve both to conserve energy and prevent disruption of the membrane permeability barrier. Our data suggest that in a comparable way the signal of non-polar residues within the N-termini of early rod/hook subunits trigger export gate opening.

## Methods

### Bacterial strains, plasmids and growth conditions

*Salmonella* strains and plasmids used in this study are listed in Table 1. The Δ*flgD*::K_m_^R^ strain in which the *flgD* gene was replaced by a kanamycin resistance cassette was constructed using the λ Red recombinase system [53]. Strains containing chromosomally encoded FliP variants were constructed by aph-I-SceI Kanamycin resistance cassette replacement using pWRG730 [54]. Recombinant proteins were expressed in *Salmonella* from the isopropyl β-D-thiogalactoside-inducible (IPTG) inducible plasmid pTrc99a [55]. Bacteria were cultured at 30–37°C in Luria-Bertani (LB) broth containing ampicillin (100 μg/ml).

**Table 1.**
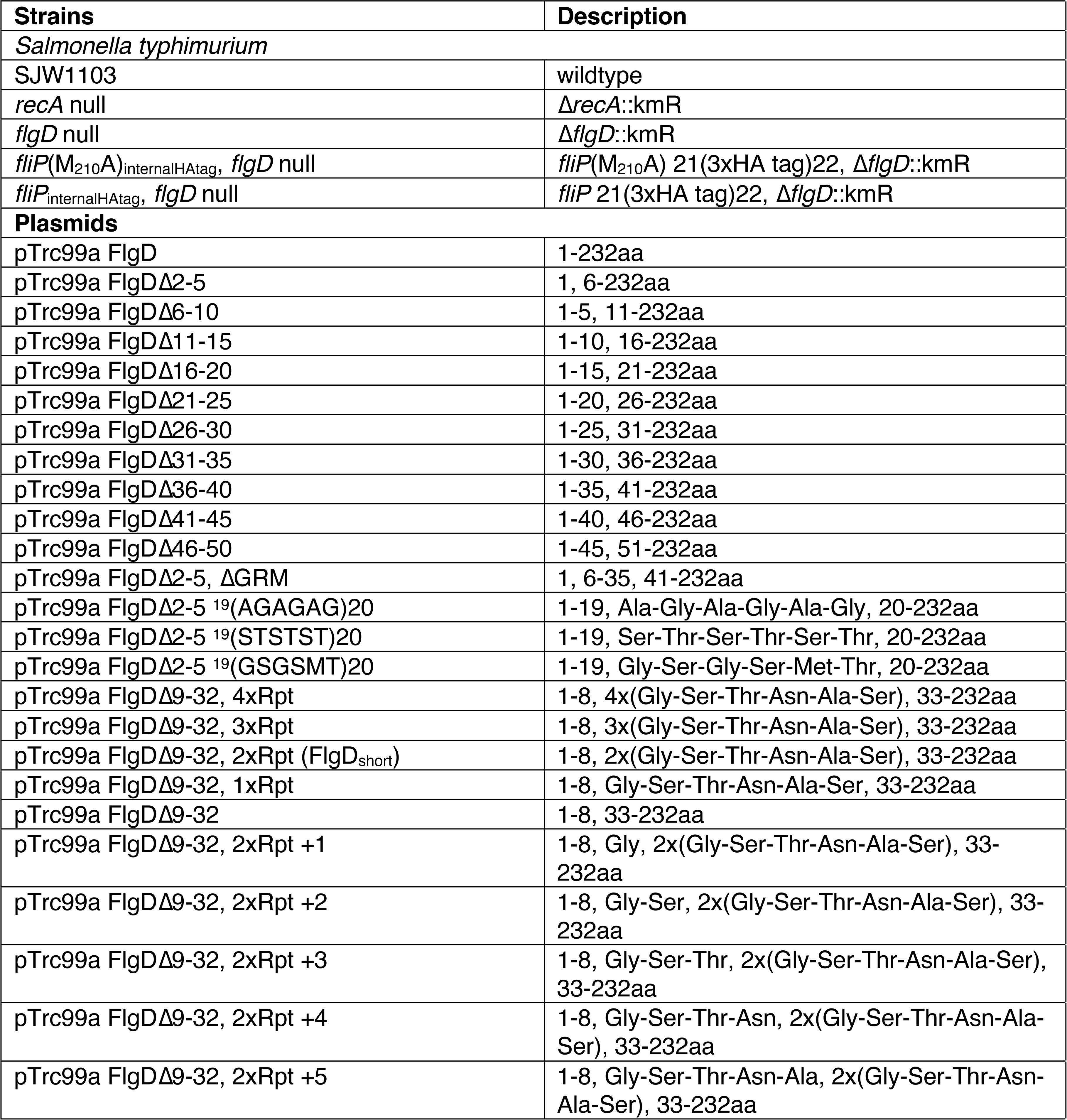
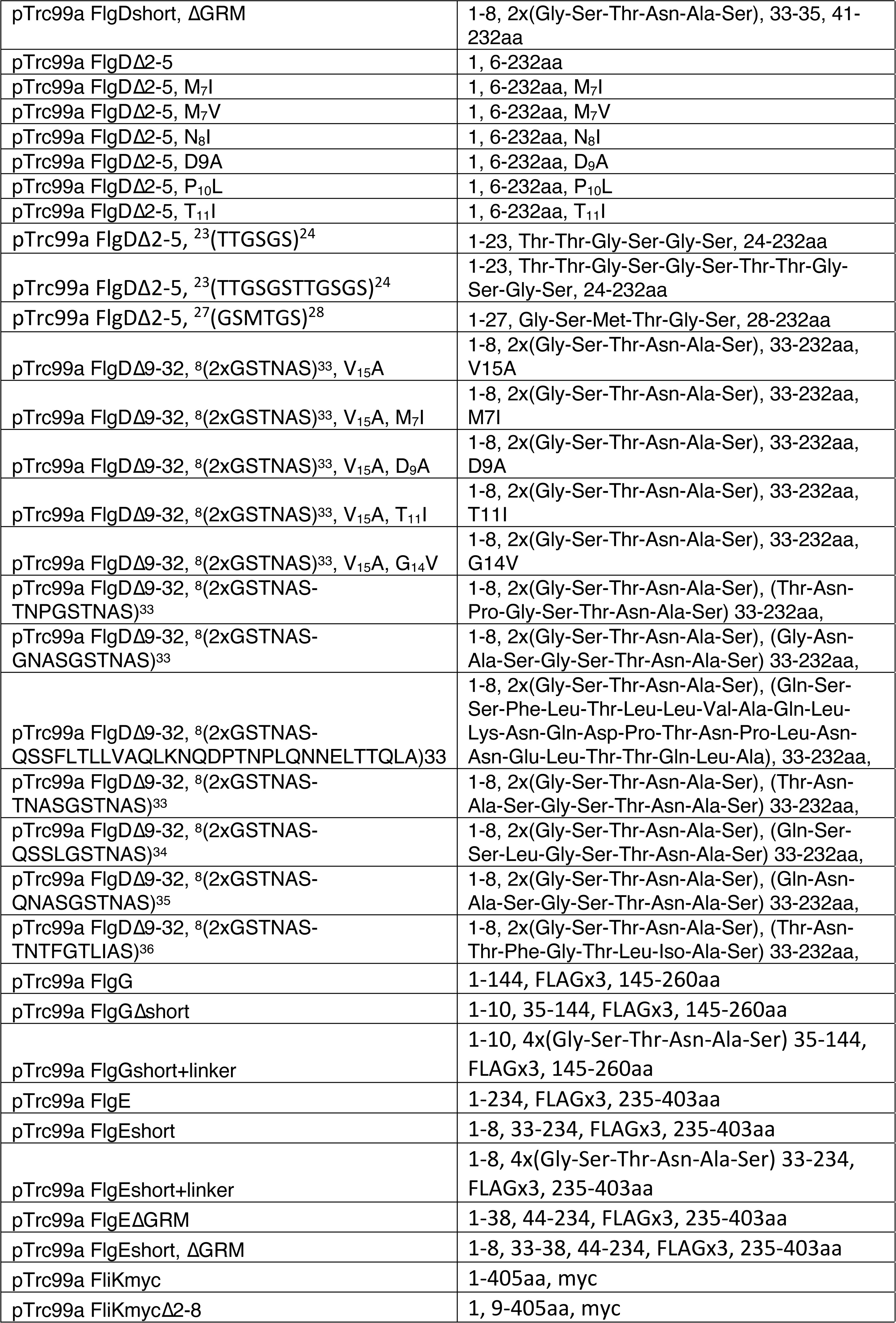
Strains and recombinant plasmids

### Flagellar subunit export assay

*Salmonella* strains were cultured at 37°C in LB broth containing ampicillin and IPTG to mid-log phase (OD_600nm_ 0.6-0.8). Cells were centrifuged (6000 x g, 3 min) and resuspended in fresh media and grown for a further 60 min at 37°C. The cells were pelleted by centrifugation (16,000 x g, 5 min) and the supernatant passed through a 0.2 μm nitrocellulose filter. Proteins were precipitated with 10% trichloroacetic acid (TCA) and 1% Triton X-100 on ice for 1 hour, pelleted by centrifugation (16,000 x g, 10 min), washed with ice-cold acetone and resuspended in SDS-PAGE loading buffer (volumes calibrated according to cell densities). Fractions were analysed by immunoblotting.

### Motility assays

For swimming motility, cultures were grown in LB broth to A600nm 1. Two microliters of culture were inoculated into soft tryptone agar (0.3% agar, 10 g/L tryptone, 5 g/L NaCl) containing ampicillin (100 μg/ml). Plates were incubated at 37°C for between 4 and 6 hours unless otherwise stated.

### Isolation of motile strains carrying suppressor mutations

Cells of the *Salmonella flgD* null strain transformed with plasmids expressing FlgD variants (FlgDΔ2-5, FlgDΔ2-5-^19^GSGSMT^20^-V_15_A or FlgD_short_) were cultured at 37°C in LB broth containing ampicillin (100 μg/ml) to mid-log phase and inoculated into soft tryptone agar (0.3% agar, 10 g/L tryptone, 5g/L NaCl) containing ampicillin (100 μg/ml). Plates were incubated at 30°C until motile ‘spurs’ appeared. Cells from the spurs were streaked to single colony and cultured to isolate the *flgD* encoding plasmid. Plasmids were transformed into the *Salmonella flgD* null strain to assess whether the plasmids were responsible for the motile suppressor phenotypes. Plasmids were sequenced to identify the suppressor mutations.

### Quantification and statistical analysis

Experiments were performed at least three times. Immunoblots were quantified using Image Studio Lite. The unpaired two-tailed Student’s *t*-test was used to determine *p*-values and significance was determined as \**p* < 0.05. Data are represented as mean ± standard error of the mean (SEM), unless otherwise specified and reported as biological replicates.

## Author contributions

**Owain J. Bryant**: Conceptualization, Data curation, Investigation, Formal analysis, Methodology, Visualisation, Writing – original draft, Writing – review & editing. **Paraminder Dhillon**: Conceptualization, Data curation, Investigation, Formal analysis, Methodology, Visualisation, Writing – review & editing. **Colin Hughes**: Conceptualization, Data curation, Formal analysis, Funding acquisition, Investigation, Methodology, Project administration, Resources, Supervision, Visualisation, Writing – review & editing. **Gillian M. Fraser**: Conceptualization, Data curation, Formal analysis, Funding acquisition, Investigation, Methodology, Project administration, Resources, Supervision, Visualisation, Writing – review & editing.

## Acknowledgements

This work was funded by grants from the Biotechnology and Biological Sciences Research Council (BB/M007197/1) to G.M.F, the Wellcome Trust (082895/Z/07/Z) to C.H. and G.M.F., a Biotechnology and Biological Sciences Research Council studentship to P.D., and a University of Cambridge John Lucas Walker studentship to O.J.B.

## Materials & Correspondence

Materials are available from the corresponding author upon request.

## Supplementary Figure Legends

**Figure S1**

Relative amounts of FlgD secreted into culture supernatants from a recombinant *Salmonella flgD* null strain expressing plasmid-based FlgDΔ2-5 suppressor mutants and wild type FlgD. Relative amounts of FlgD secreted into culture supernatants were quantified using image Studio Lite software and normalised to the level of wild type FlgD in culture supernatants. Error bars represent the standard error of the mean calculated from at least three biological replicates.

**Figure S2.**

**a.** N-terminal sequences of all *Salmonella* flagellar rod and hook subunits aligned to their gate-recognition motif (GRM, blue). Small non-polar residues upstream from the gate-recognition motif are highlighted (yellow).

**b.** Hydrophobicity plots for the N-terminal 60 residues of each *Salmonella* flagellar rod and hook subunit were generated by ExPASy tools using the Kyte and Doolittle method [56]. The *x* axis of the plot indicates the amino acid position, starting from the N terminus. The *y* axis of the plot indicates the hydrophobicity of the amino acid sequence, where higher values represent higher hydrophobicity. Amino acid sequence corresponding to the gate-recognition motif of each subunit is highlighted in blue.

**Figure S3**

Whole cell (cell) and supernatant (secreted) proteins from late exponential-phase cultures of *Salmonella flgD* null strains expressing wild type FliK containing a C-terminal myc tag for immunodetection (FliK) or its variant in which residues two to eight are deleted (FliKΔ2-8) were separated by SDS (15%)-PAGE and analysed by immunoblotting with anti-myc monoclonal antisera (top), anti-FlhA (middle) or anti-FlgN (bottom) polyclonal anti-sera. Apparent molecular weights are in kilodaltons (kDa).

**Figure S4**

**a.** Swimming motility of a *Salmonella* Δ*recA* strain expressing: suppressor mutants isolated from the FlgDΔ2-5 variant (labelled as FlgDΔ2-5 N_8_I or FlgDΔ2-5 T_11_I), the parent FlgDΔ2-5 variant (labelled as DΔ2-5) or wild type FlgD (labelled as FlgD). Motility was assessed in 0.25% soft-tryptone agar containing 100 μg/ml ampicillin and 50 μM IPTG and incubated 4-6 hours at 37°C.

**b.** Swimming motility of a *Salmonella* Δ*recA* strain expressing wild type FlgD (labelled as FlgD), FlgDΔ2-5 (labelled as DΔ2-5) or its variants containing a six-residue insertion between residues 19 and 20 of either small non-polar (AGAGAG) residues (labelled as DΔ2-5 3x(AG)), polar (STSTST) residues (labelled as DΔ2-5 3x(ST)), or the sequence from an isolated insertion suppressor mutant (GSGSMT) (labelled as DΔ2-5 GSGSMT). Motility was assessed in 0.25% soft-tryptone agar containing 100 μg/ml ampicillin and 50 μM IPTG and incubated 4-6 hours at 37°C.

**Figure S5**

**a.** Whole cell (cell) and supernatant (secreted) proteins from late exponential-phase cultures of *Salmonella flgD* null strains expressing: suppressor mutants isolated from the FlgDΔ2-5 variant (FlgDΔ2-5 N_8_I or FlgDΔ2-5 T_11_I), FlgDΔ2-5 variant (-) or wild type FlgD (FlgD) were separated by SDS (15%)-PAGE and analysed by immunoblotting with anti-FlhA or FlgN polyclonal antisera. Apparent molecular weights are in kilodaltons (kDa).

**b.** Whole cell (cell) and supernatant (secreted) proteins from late exponential-phase cultures of *Salmonella flgD* null strains expressing wild type FlgD (FlgD), FlgDΔ2-5 (labelled as DΔ2-5) or its variants containing a six-residue insertion between residues 19 and 20 of either small non-polar (AGAGAG) residues (labelled as DΔ2-5 3x(AG)), polar (STSTST) residues (labelled as DΔ2-5 3x(ST)), or the sequence from an isolated insertion suppressor mutant (GSGSMT) (labelled as DΔ2-5 GSGSMT) were separated by SDS (15%)-PAGE and analysed by immunoblotting with anti-FlgD polyclonal antisera. Apparent molecular weights are in kilodaltons (kDa).

**Figure S6**

**a.** Whole cell (cell) and supernatant (secreted) proteins from late exponential-phase cultures of a *Salmonella flgD* null strain expressing suppressor mutants isolated from a FlgDΔ2-5-^19^(GSGSMT)^20^-V_15_A variant (V_15_A-M_7_I, V_15_A-D_9_A, V_15_A-T_11_I, V_15_A-G_14_V), their parent FlgD variant FlgDΔ2-5-^19^(GSGSMT)^20^-V_15_A (labelled as V_15_A), FlgDΔ2-5-^19^(GSGSMT)^20^ (-) or wild type FlgD (FlgD) were separated by SDS (15%)-PAGE and analysed by immunoblotting with anti-FlhA and FlgN polyclonal antisera. Apparent molecular weights are in kilodaltons (kDa).

**b.** Whole cell (cell) and supernatant (secreted) proteins from late exponential-phase cultures of a *Salmonella flgD* null strain expressing wild type FlgD (FlgD), FlgDΔ9-32 or its variants in which residues 9-32 were replaced by between one and four six-residue repeats of Gly-Ser-Thr-Asn-Ala-Ser (GSTNAS): (Δ9-32 4xRpt, Δ9-32 3xRpt, Δ9-32 2xRpt, Δ9-32 1xRpt) were separated by SDS (15%)-PAGE and analysed by immunoblotting with anti-FlhA and anti-FlgN polyclonal antisera. Apparent molecular weights are in kilodaltons (kDa).

**c.** Whole cell (cell) and supernatant (secreted) proteins from late exponential-phase cultures of a *Salmonella flgD* null strain expressing wild type FlgD (labelled as FlgD), a FlgD variant in which residues 9-32 were replaced by two repeats of a six-residue sequence Gly-Ser-Thr-Asn-Ala-Ser (labelled as 2xRpt) or its variants containing between one and five additional residues inserted directly after the two repeats (labelled as 2xRpt+ 1, 2xRpt+ 2, 2xRpt+ 3, 2xRpt+ 4 or 2xRpt+ 5) were separated by SDS (15%)-PAGE and analysed by immunoblotting with anti-FlgD polyclonal antisera. Apparent molecular weights are in kilodaltons (kDa).

**d.** Whole cell (cell) and supernatant (secreted) proteins from late exponential-phase cultures of a *Salmonella flgD* null strain expressing wild type FlgD (labelled as FlgD), a FlgD variant in which residues 9-32 were replaced by two repeats of a six-residue sequence Gly-Ser-Thr-Asn-Ala-Ser (labelled as FlgD_short_) or suppressor mutants isolated from this strain (labelled as rev1, rev2 or rev3) were separated by SDS (15%)-PAGE and analysed by immunoblotting with anti-FlhA and anti-FlgN polyclonal antisera. Apparent molecular weights are in kilodaltons (kDa).

**Figure S7**

**a.** Whole cell (cell) and supernatant (sec) proteins from late exponential-phase cultures of a *Salmonella flgE* null strain expressing wild type FlgG (labelled as FlgG_wild type_), a FlgG variant in which residues 11-35 were deleted (labelled as FlgG_short_) or a FlgG variant in which residues 11-35 were replaced by four repeats of a six-residue sequence Gly-Ser-Thr-Asn-Ala-Ser (labelled as FlgG_short_+4Rpt). All FlgG variants were engineered to contain an internal 3xFLAG tag for immunodetection. Proteins were separated by SDS (15%)-PAGE and analysed by immunoblotting with anti-FlhA or anti-FlgN polyclonal antisera. Apparent molecular weights are in kilodaltons (kDa).

**b.** Whole cell (cell) and supernatant (secreted) proteins from late exponential-phase cultures of a *Salmonella flgD* null strain expressing wild type FlgE (labelled as FlgE_wild type_), a FlgE variant in which residues 9-32 were deleted (labelled as FlgE_short_) or a FlgE variant in which residues 9-32 were replaced by four repeats of a six-residue sequence Gly-Ser-Thr-Asn-Ala-Ser (labelled as FlgE_short_+4Rpt). All FlgE variants were engineered to contain an internal 3xFLAG tag for immunodetection. Proteins were separated by SDS (15%)-PAGE and analysed by immunoblotting with anti-FLAG monoclonal antisera.

**Figure S8**

**a.** Whole cell (cell) proteins from late exponential-phase cultures of recombinant *Salmonella flgD* null strains producing a chromosomally-encoded FliP-M_210_A variant (M_210_A gate, left) or wild type FliP (wild type gate, right). Wild type FliP and FliP-M_210_A were engineered to contain an internal HA tag positioned between residue 21 and 22 to allow immunodetection of FliP [30] (bottom panel). Both strains produced either a pTrc99a plasmid-encoded FlgD subunit variant in which residues 9-32 were replaced with two repeats of the six amino acid sequence Gly-Ser-Thr-Asn-Ala-Ser (FlgD_short_; top panel) or a pTrc99a plasmid-encoded wild type FlgD subunit (FlgD_wild type_; middle panel). Proteins were separated by SDS (15%)-PAGE and analysed by immunoblotting with anti-FlgD polyclonal antisera or anti-HA monoclonal antisera.

**b.** The mean motility halo diameter of recombinant *Salmonella flgD* null strains producing a chromosomally-encoded FliP-M_210_A variant (M_210_A gate, left) or wild type FliP (wild type gate, right). Wild type FliP and FliP-M_210_A were engineered to contain an internal HA tag positioned between residue 21 and 22 to allow immunodetection of FliP. Both strains produced a pTrc99a plasmid-encoded wild type FlgD. Error bars represent the standard error of the mean calculated from at least three biological replicates. *** indicates a p-value <0.001.

**Figure S9**

**a.** The N-terminal sequence of the *Salmonella* FlgD cap subunit aligned by the conserved gate-recognition motif (GRM, blue) with N-terminal sequences of FlgD from other bacterial species. Small non-polar residues located N-terminal from the minimum distance threshold of 24 residues from the gate-recognition motif are highlighted (yellow).

**b.** The N-terminal sequence of the *Salmonella* FlgE hook subunit aligned by the conserved gate-recognition motif (GRM, blue) with N-terminal sequences of FlgE from other bacterial species. Small non-polar residues located N-terminal from the minimum distance threshold of 24 residues from the gate-recognition motif are highlighted (yellow).

**c.** The N-terminal sequence of the *Salmonella* FliK ruler subunit aligned by the conserved gate-recognition motif (GRM, blue) with N-terminal sequences of FliK from other bacterial species. Small non-polar residues located N-terminal from the minimum distance threshold of 24 residues from the gate-recognition motif are highlighted (yellow).

**Figure S10**

**a.** Cryo-EM structure of the FliPQR-FlhB complex (PDB:6S3L [31], left) displaying FliPQR (green) and FlhB residues 1-221 (red), the alphafold predicted structure of full length FlhB (red, middle) and the Cryo-EM structure of the PrgHK-SpaPQR complex (PDB:6PEM) [57] displaying PrgHK (magneta) in complex with the FliPQR homologue, SpaPQR (green).

**b.** Structures from (a) were superimposed in chimera using MatchMaker to generate a predicted model of the PrgHK - FliPQR - full length FlhB complex (left). A tomographic reconstruction of the *Salmonella* SPI-1 vT3SS (EMD-20838) [28] is shown in grey (right).

**c.** Placement of the PrgHK – FliPQR – full length FlhB predicted model in the tomographic reconstruction of the Salmonella SPI-1 vT3SS, which suggests that the substrate binding cytoplasmic domain of FlhB (FlhB_C_) is positioned below the plane of inner membrane and above the visible nonameric ring formed by the cytoplasmic domain of FlhA (FlhA_C_). Using the predicted model, the distance between the GRM binding site on FlhB_C_ and the base of the FliPQR-FlhB_N_ complex is estimated to be between 70 and 80 angstrom.

**Figure S11**

A schematic of an early flagellar subunit containing an N-terminal export signal (yellow) and gate-recognition motif (GRM, blue). The essential gate recognition motif (GRM) of *Salmonella* FlgD (residues 36-40) was aligned with homologous regions in other early flagellar subunits and with the inner rod and ruler subunits of the injectisome from *Salmonella* and other bacterial species.

