## Supplementary figures and images for "Recognition of discrete export signals in early flagellar subunits during bacterial Type III secretion"

### Figure S1

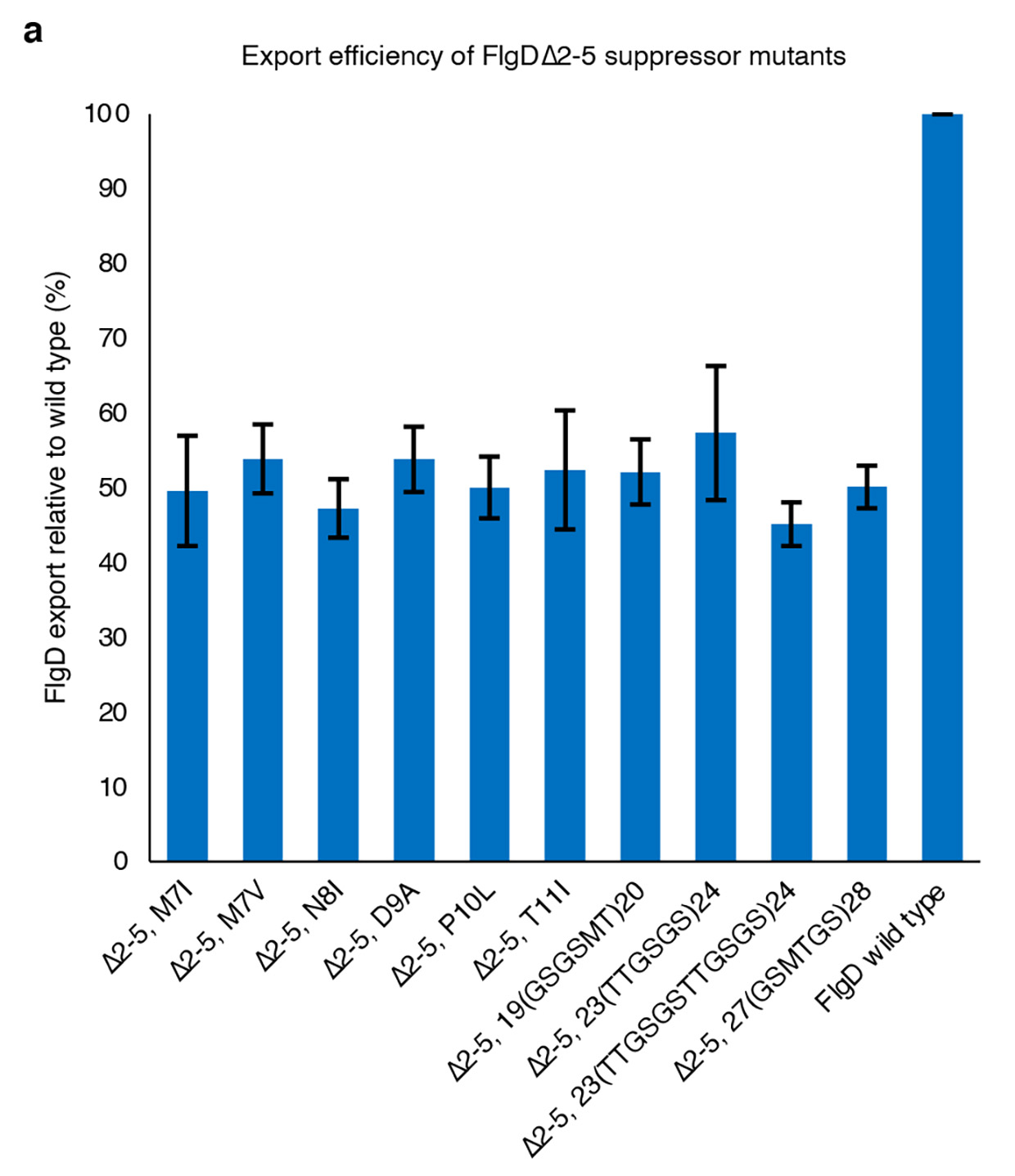

### Figure S2

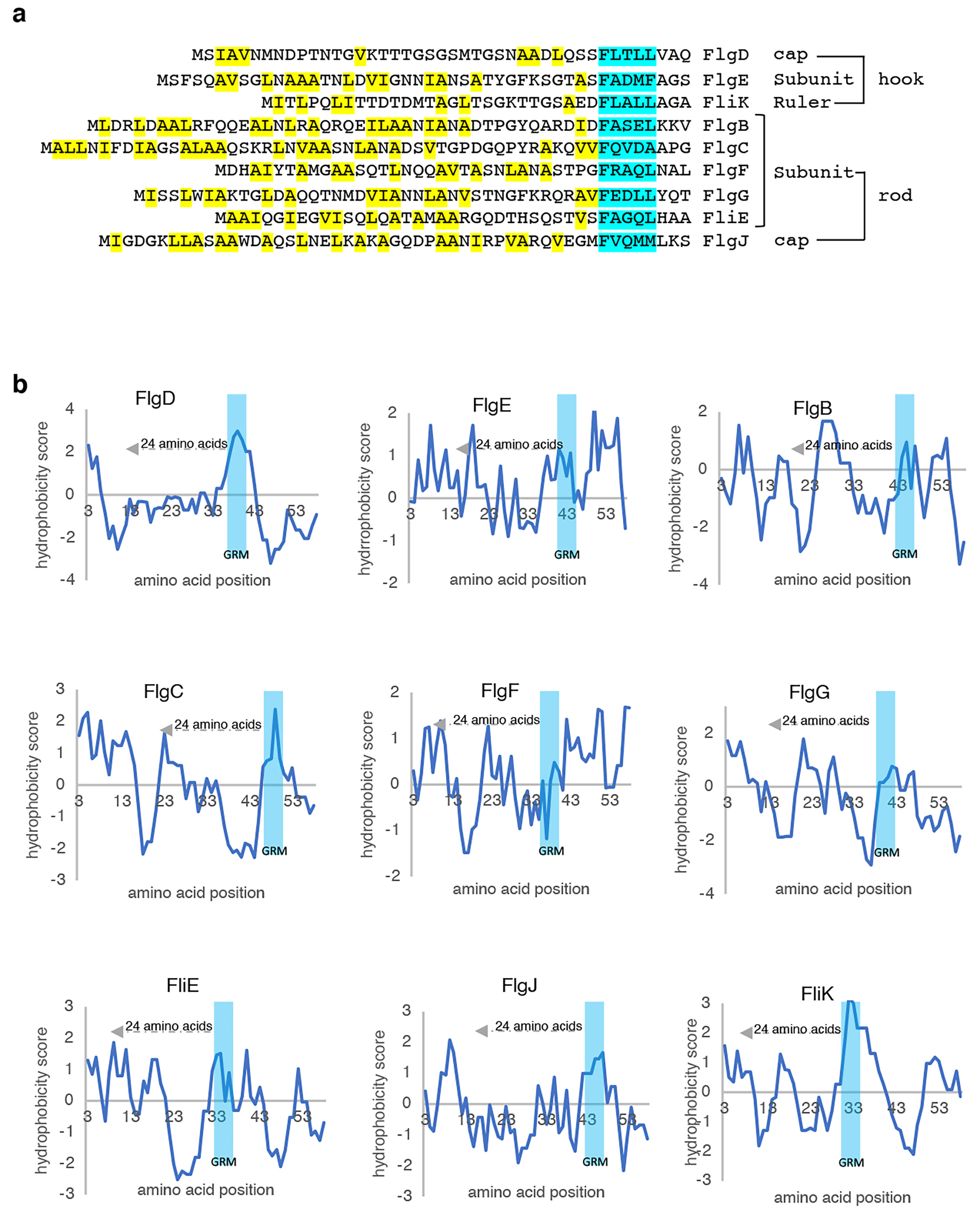

### Figure S3

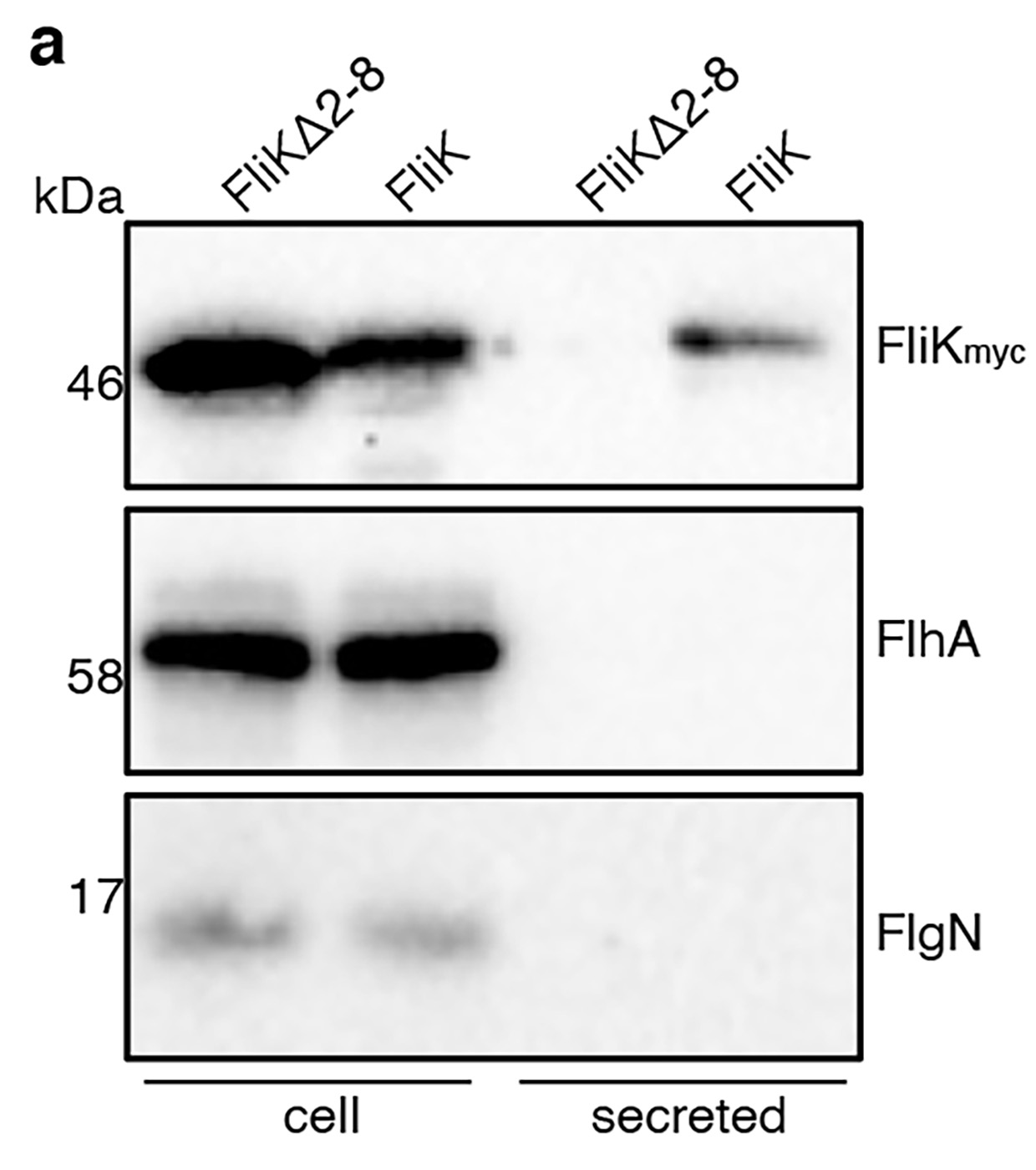

### Figure S4

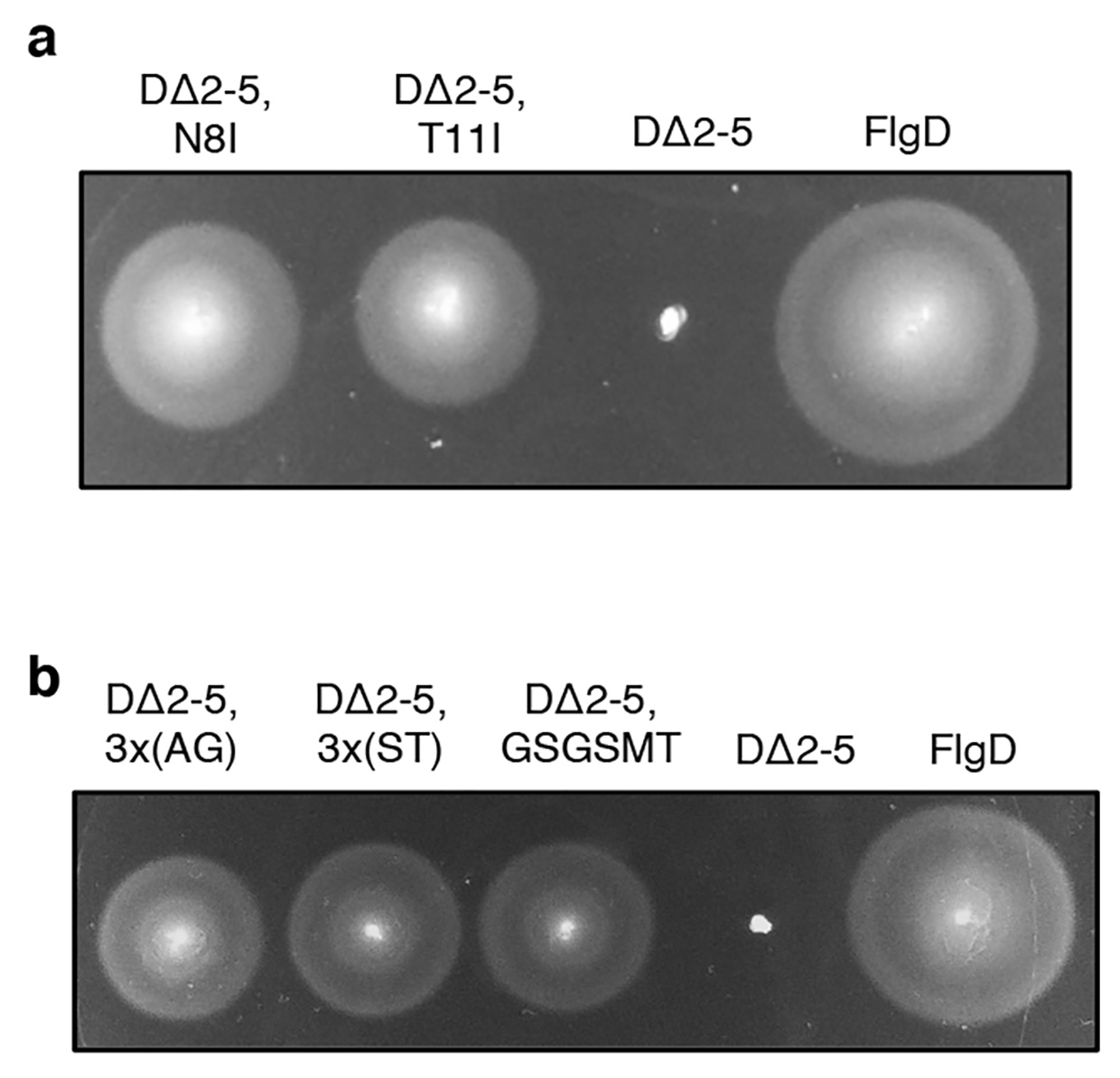

### Figure S5

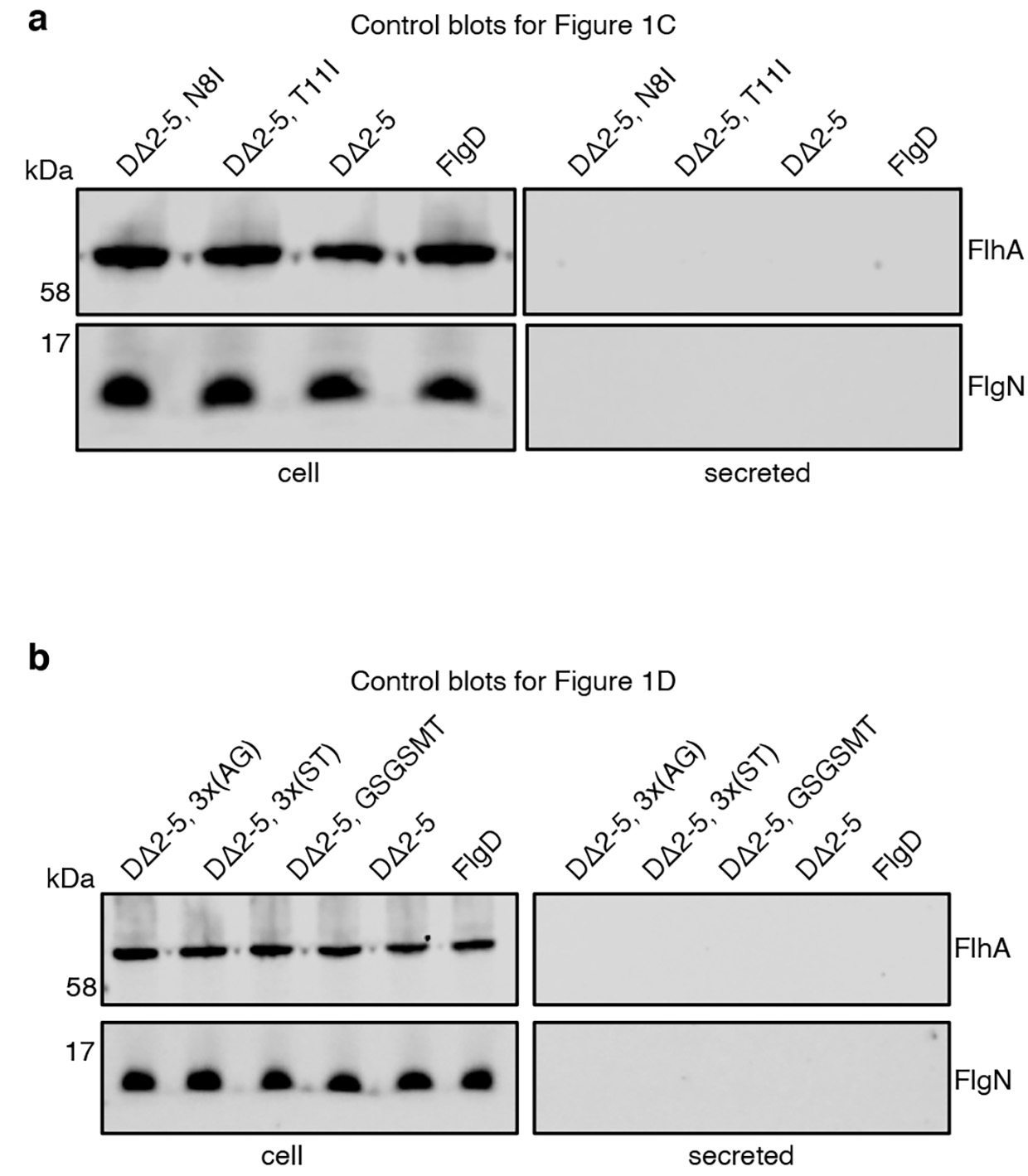

### Figure S6

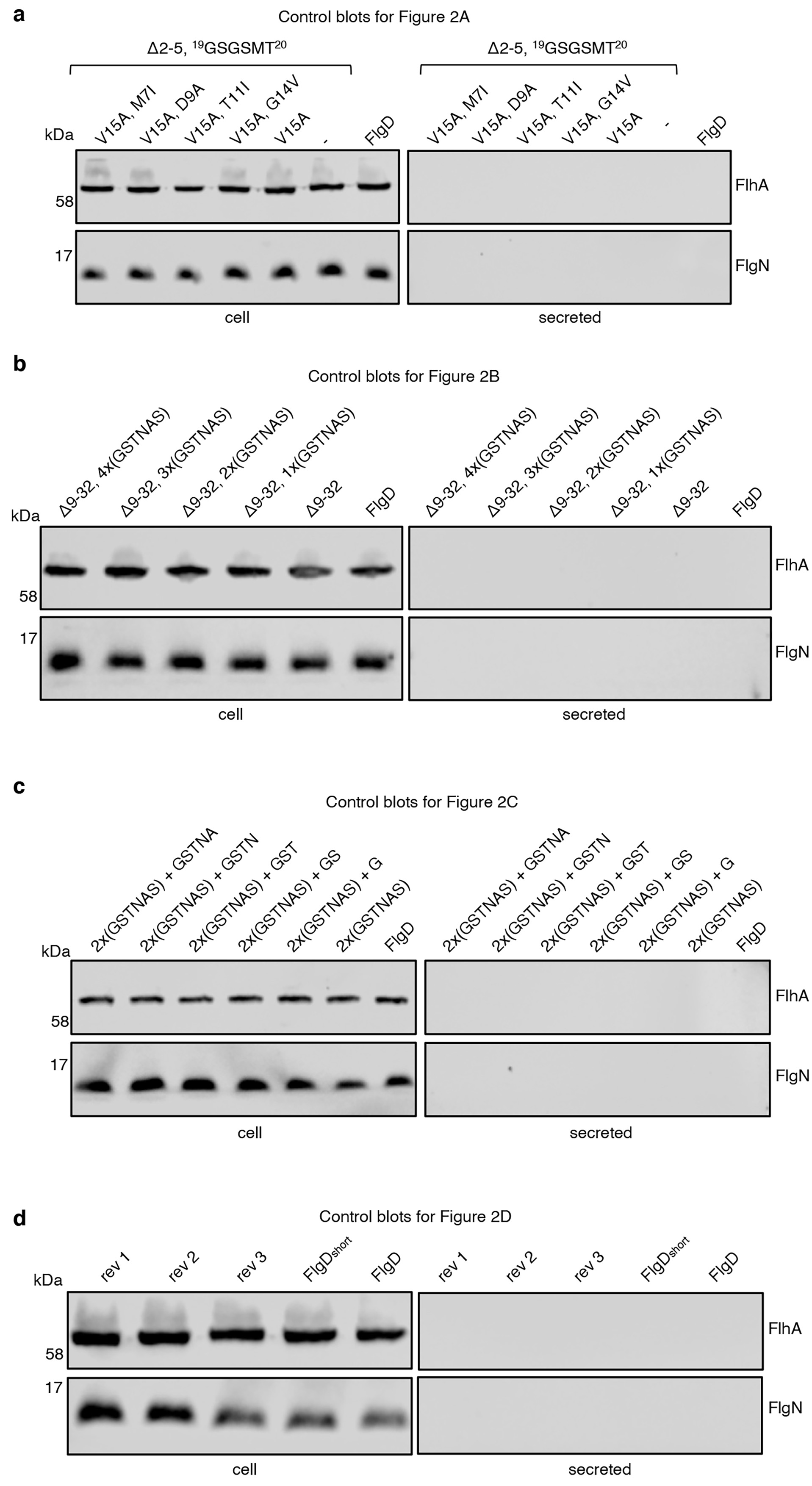

### Figure S7

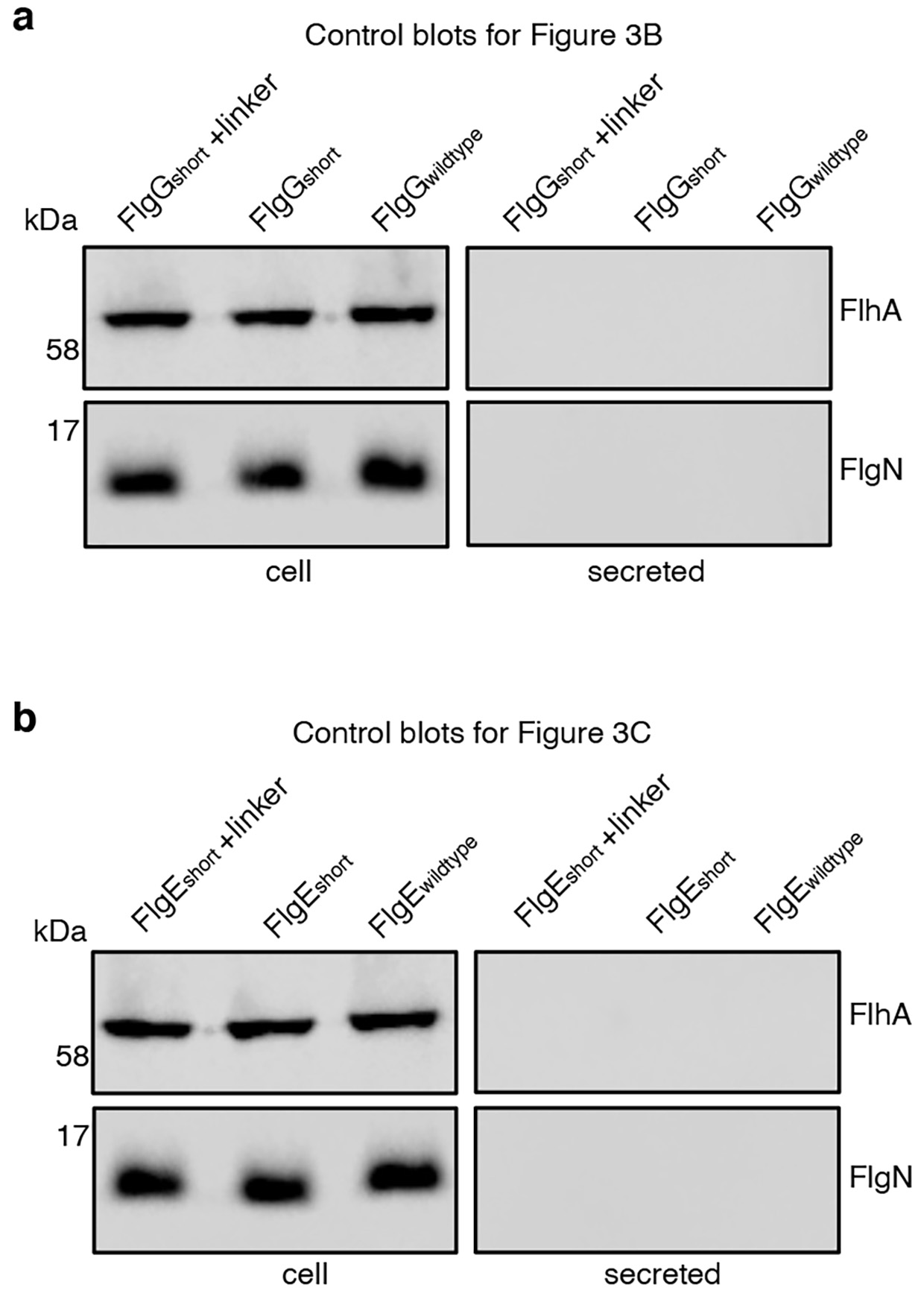

### Figure S8

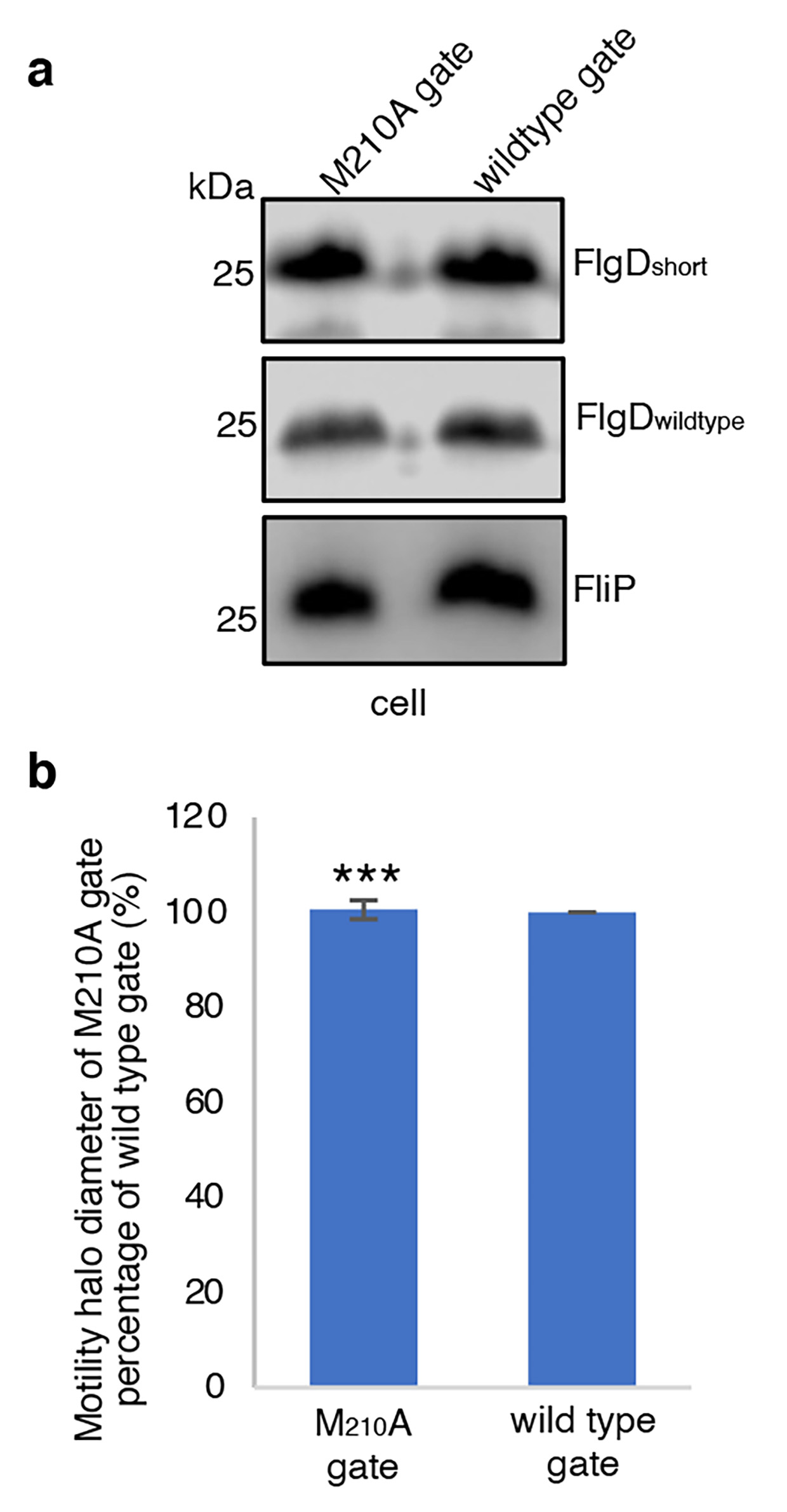

### Figure S9

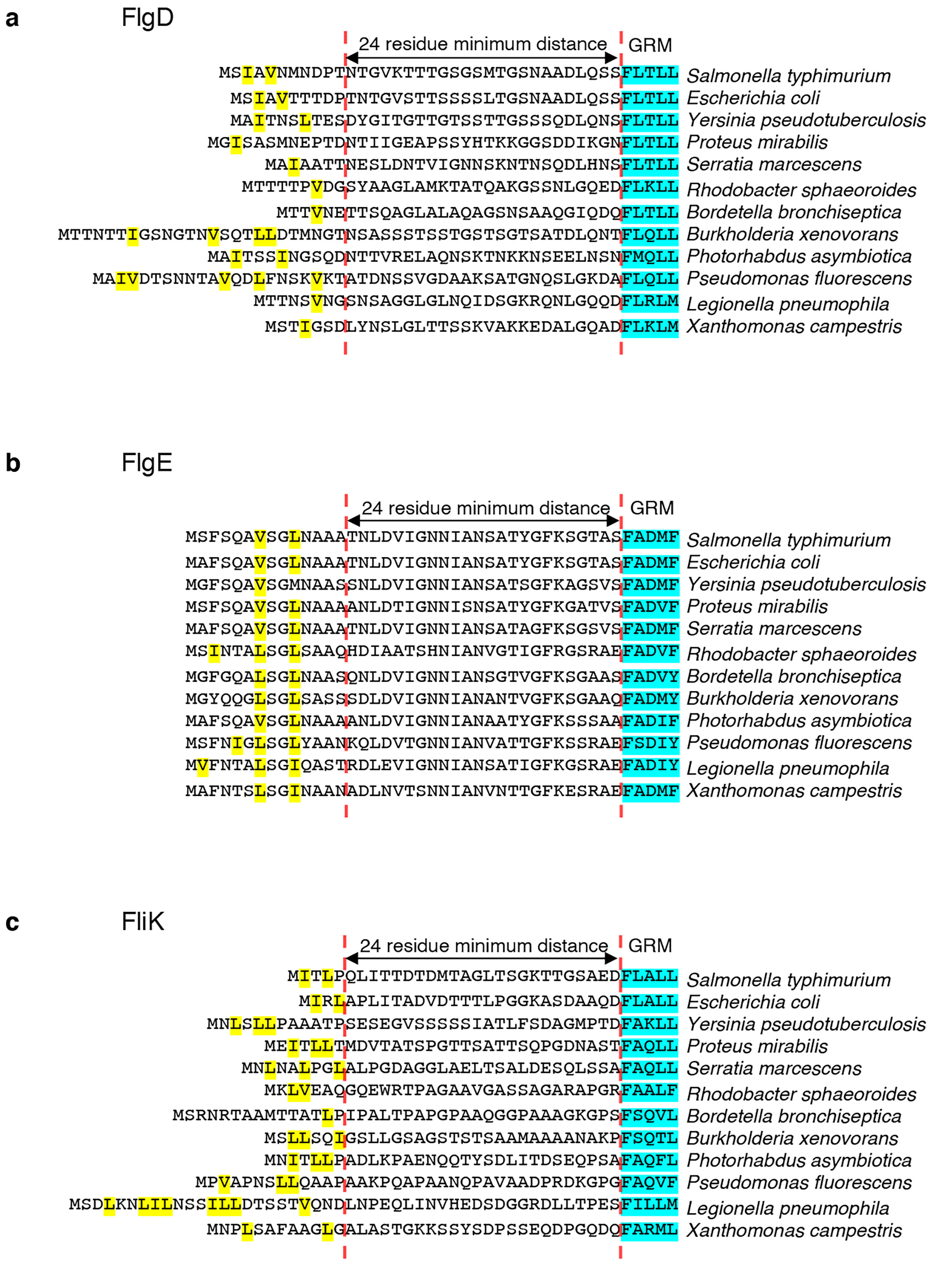

### Figure S10

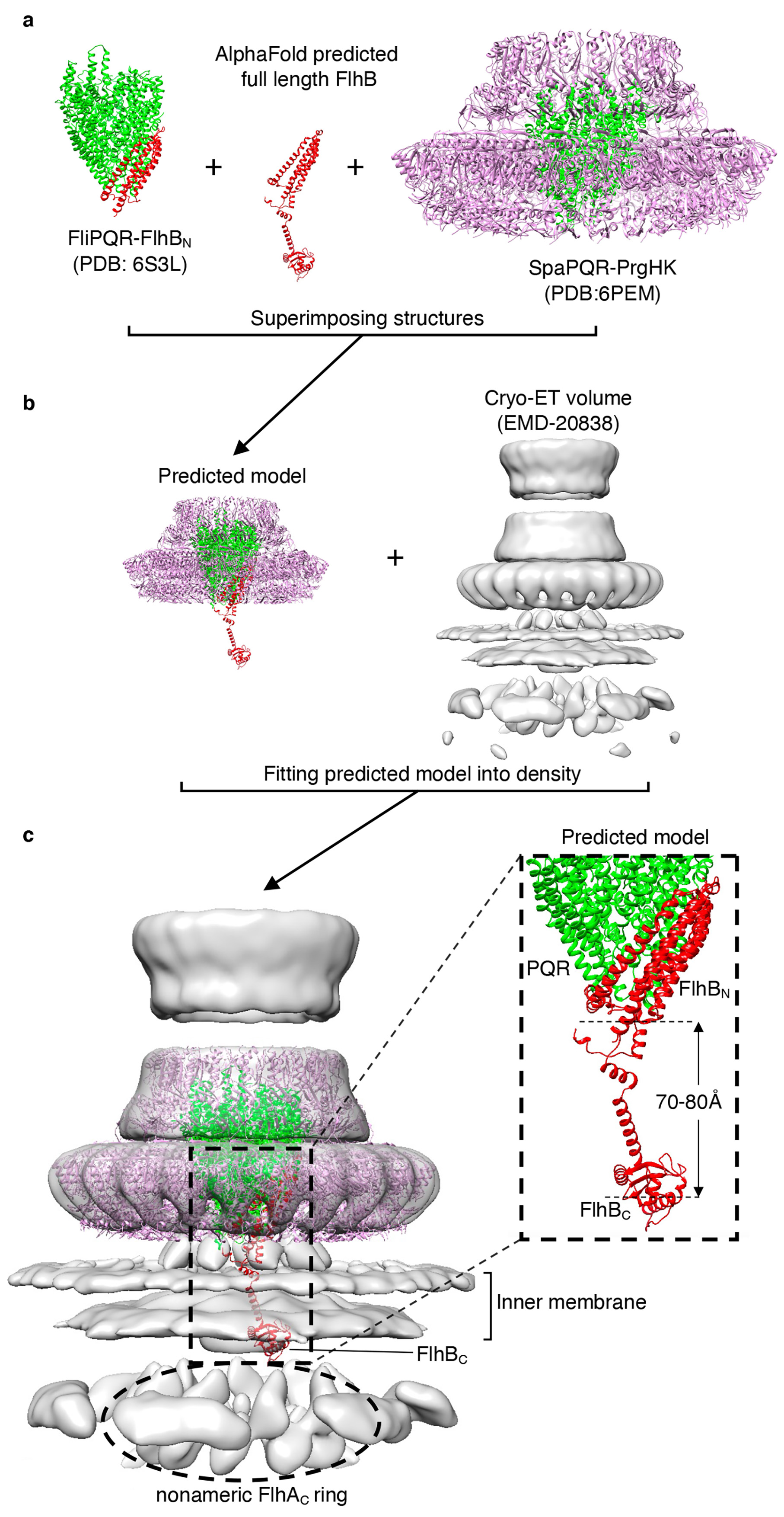

### Figure S11

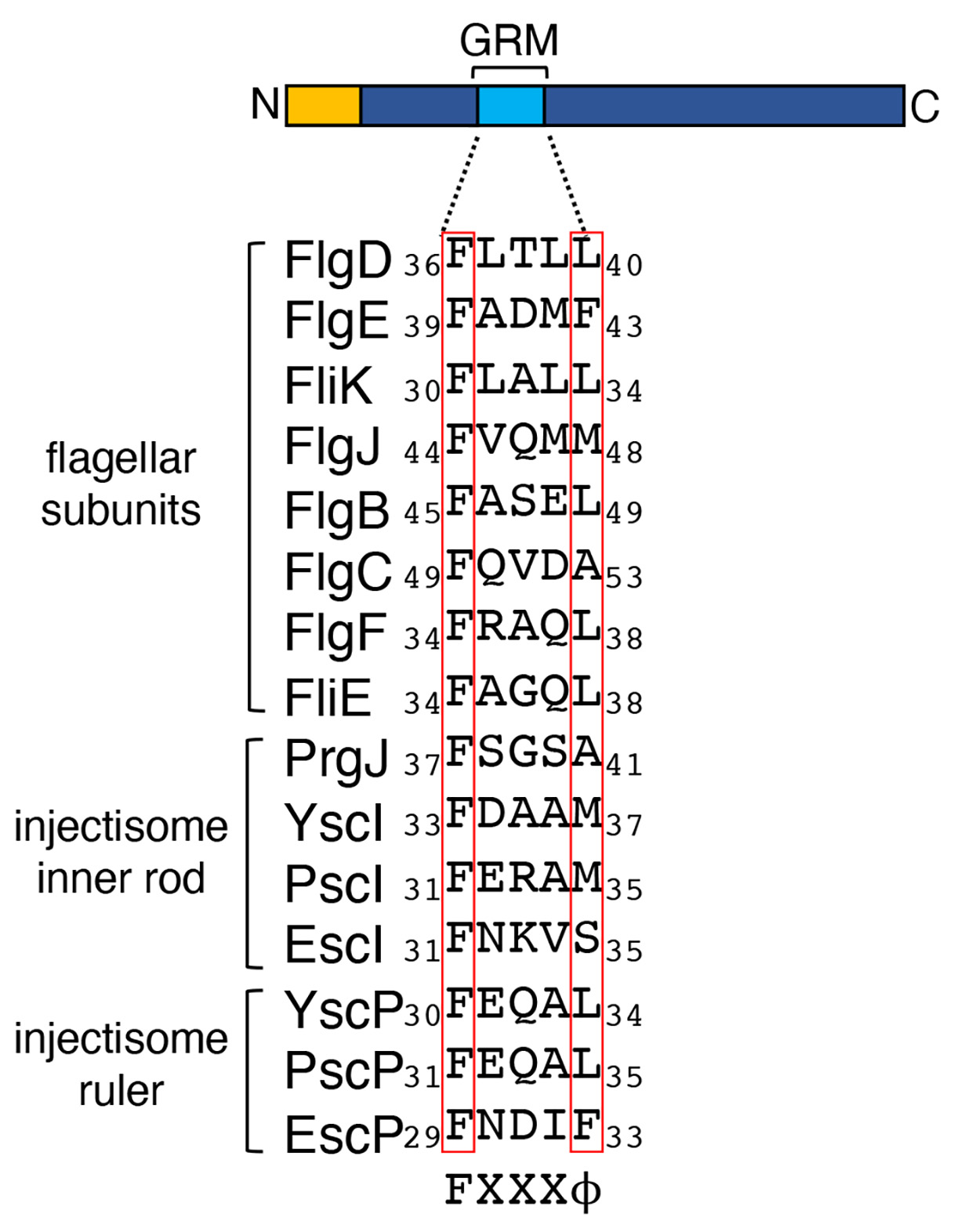
